# Breathing coordinates limbic network dynamics underlying memory consolidation

**DOI:** 10.1101/392530

**Authors:** Nikolaos Karalis, Anton Sirota

**Author notes:** Present address: Friedrich Miescher Institute for Biomedical Research, 4058, Basel, Switzerland (Nikolaos Karalis). Correspondence (N.K.), (A.S.).

## Abstract

The coordinated activity between remote brain regions underlies cognition and memory function. Although neuronal oscillations have been proposed as a mechanistic substrate for the coordination of information transfer and memory consolidation during sleep, little is known about the mechanisms that support the widespread synchronization of brain regions and the relationship of neuronal dynamics with other bodily rhythms, such as breathing. Here we address this question using large-scale recordings from a number of structures, including the medial prefrontal cortex, hippocampus, thalamus, amygdala and nucleus accumbens in mice. We identify a dual mechanism of respiratory entrainment, in the form of an intracerebral corollary discharge that acts jointly with an olfactory reafference to coordinate limbic network dynamics, such as hippocampal ripples and cortical UP and DOWN states, involved in memory consolidation. These results highlight breathing, a perennial rhythmic input to the brain, as an oscillatory scaffold for the functional coordination of the limbic circuit, enabling the segregation and integration of information flow across neuronal networks.

---

Over the past century, cortical and subcortical structures of the limbic circuit and the medial temporal lobe have been identified as critical elements of the memory circuit, involved in emotional regulation and the formation, consolidation, and retrieval of episodic memories^1–3^. Although the anatomical substrate of these circuits has been elaborated in detail, mechanisms that enable the processing and transfer of information across these distributed circuits are not well understood.

Neuronal dynamics are characterized by oscillatory activity associated with distinct behavioral states and functional roles^4^. During active states, hippocampal theta oscillations dynamically coordinate local activity and information flow between the hippocampus and entorhinal cortex^5–7^, as well as other limbic structures such as the medial prefrontal cortex (mPFC)^8,9^.

During slow-wave sleep, the cortex is in a bistable state, characterized by spontaneous alternations between UP and DOWN states in the membrane potential and action potential firing of neurons^10,11^. In parallel, the hippocampal neurons are engaged in transient, fast oscillatory events termed sharp-wave ripples (SWR), during which awake activity is replayed^12^. Such nonlinear dynamics are coordinated between regions^13–15^ and their interaction is believed to support memory consolidation^16,17^ and the transfer of memories to their permanent cortical storage^18,19^.

While the importance and role of the cortical slow oscillation (SO), hippocampal ripples, and their coordination during sleep have been established, the mechanisms that support this coordination across distributed cortical and limbic circuits during sleep remain elusive and a global pacemaker that ties together distinct network dynamics has not been identified. Recently, a number of studies have identified signatures of respiration in the cortical and hippocampal LFP of rodents^20–23^ and humans ^24,25^, which has been attributed to reafferent olfactory activity. However, the function of these phenomena is not understood and the mechanism and consequences of respiratory modulation have not been established, given the limitations in interpreting LFP signals.

Here we address the hypothesis that the breathing rhythm modulates the activity of the limbic brain and underlies the coordination of network dynamics across limbic systems during offline states. To this end, using high-density silicon probes we performed a large-scale *in vivo* functional anatomical characterization of the medial prefrontal cortex (mPFC), hippocampus, basolateral amygdala (BLA) and nucleus accumbens (NAc). Using this approach, we identified an intracerebral centrifugal respiratory corollary discharge that acts in concert with a respiratory reafference and mediates the inter-regional synchronization of limbic memory circuits. The respiratory modulation is acting as a functional oscillatory scaffold, that together with the underlying anatomical substrate, organizes information flow and systems memory consolidation processes.

## Results

### Respiratory entrainment of prefrontal cortex across brain states

To investigate the role of breathing in organizing neuronal dynamics in the medial prefrontal cortex (mPFC), we recorded simultaneously the local electrical activity (electroolfactogram; EOG)^26^ of the olfactory sensory neurons (OSNs) and singleunit and LFP in the mPFC in freely-behaving mice (**Fig. 1a**). The EOG reflected the respiratory activity and exhibited reliable phase relationship to the respiratory cycle, as established by comparing this signal to the airflow from the nostrils (Supplementary Fig. 1a-d), and was reflected in rhythmic head-motion (Supplementary Fig. 1b). We then segmented behavioral states based on the head micro-motion (Supplementary Fig. 1e-f), differentiating slow wave sleep (SWS), REM sleep, quiescence and awake exploration. Behavioral state changes were associated with changes in the breathing frequency (**Fig. 1c, Supplementary Fig.2a**) which are accompanied by autonomic changes conferred upon the heart by central regulation and respiratory sinus arrhythmia (RSA), indicative of a generalized role of breathing in coordinating bodily rhythms (**Supplementary Fig. 2b-f**).

**Figure 1.**
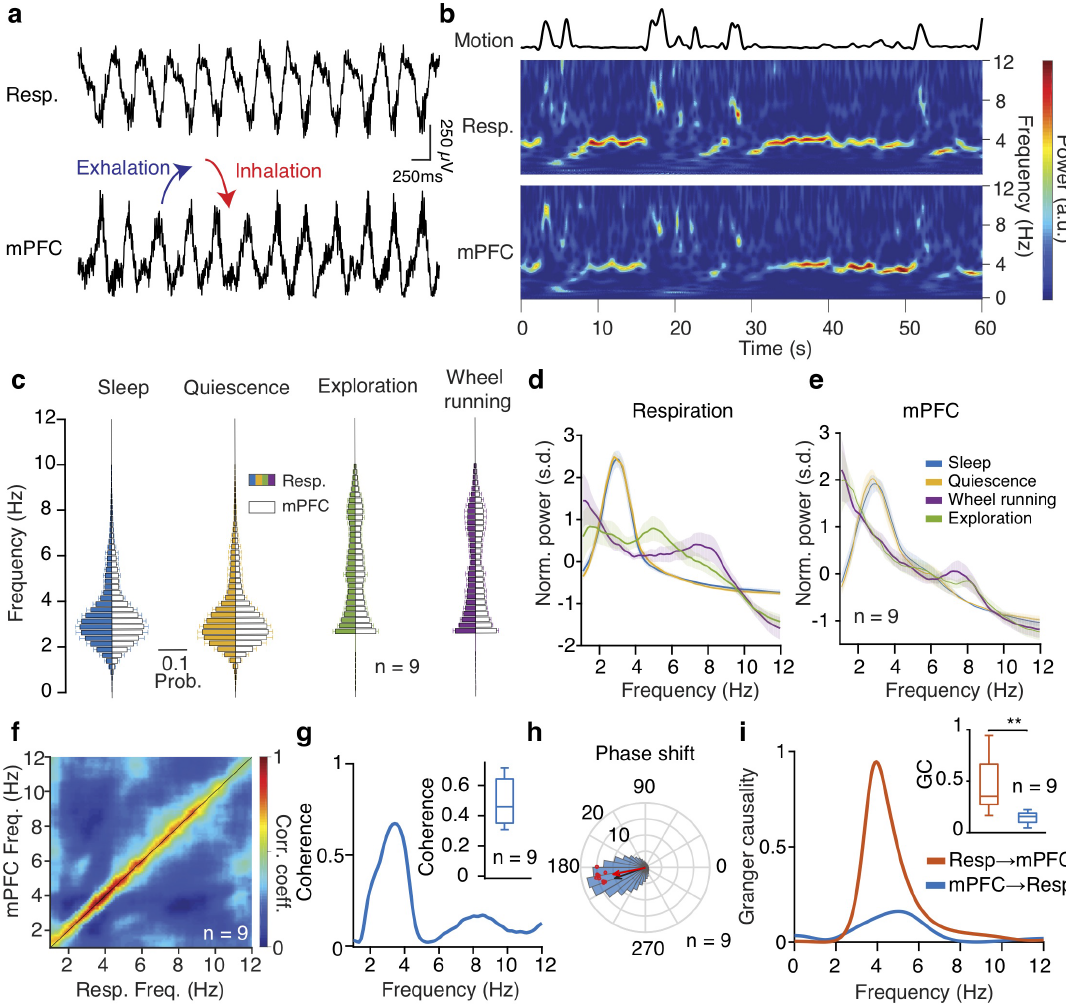
Prefrontal oscillations are related to respiration throughout behavioral states. (**a**) Example traces of simultaneously recorded respiratory EOG and medial prefrontal local field potentials (LFP). (**b**) Example time-frequency decomposition of respiratory and mPFC LFP signals, revealing the reliable relationship between the two signals. (**c**) Distribution of peak frequency bins of the spectrally decomposed respiration (left; darker colors) and mPFC LFP (right; lighter colors) during slow-wave sleep, quiescence, exploratory behavior and self-initiated wheel running (n = 9 mice). (**d**, **e**) Averaged normalized power spectral density of respiration (**d**) and mPFC LFP (**e**) across states as in (**c**). (**f**) Frequency-resolved comodulation of respiration and mPFC LFP oscillation power, across mice and behaviors (n = 9 mice). (**g**) Example coherence spectrum between respiration and mPFC LFP during offline states. Inset, average coherence value in the 2-5 Hz band (n = 9 mice). (**h**) Phase shift of 2-5 Hz filtered respiration and mPFC LFP signals during offline states for an example animal (blue histogram) and overlaid magnitude of phase modulation (logZ) and average phase shift for all animals (red dots; n = 9 mice). Black arrow depicts the average phase and logZ of the phase shift for the example and the red arrow for the population. (**i**) Example spectral Granger causality between respiration and mPFC LFP for both causal directions. Inset, average Granger causality for the 4 Hz band (2–5 Hz) between respiration and mPFC LFP for both causality directions (n = 9 mice, Wilcoxon signed-rank test, resp → mPFC versus mPFC → resp, ** P<0.01). a.u., arbitrary units; s.d., standard deviations. Shaded areas, mean ± s.e.m.

Examination of the spectrotemporal characteristics of respiratory activity and the prefrontal LFP revealed a faithful reflection of the respiratory activity in the prefrontal LFP (**Fig. 1b**), suggesting a potential relationship between these oscillations. The two oscillations were comodulated across a wide frequency range (**Fig. 1f**) and this relationship was preserved throughout many active (online) as well as inactive (offline) states in freely-behaving mice (**Fig. 1c-e**). Coherence and Granger causality analysis of the respiratory and LFP signals suggested that the respiratory and LFP signals suggested that the respiratory oscillation is tightly locked and likely causally involved in the generation of the prefrontal LFP oscillation signal (**Fig. 1g-i**). During fear behavior, the mouse mPFC is dominated by a prominent 4 Hz oscillation^27^. The similarity in frequency suggests that respiration is the origin of fear-related 4 Hz oscillations, To explicitly test this, we exposed mice to auditory, contextual and innate fear paradigms (Supplementary Fig. 3a,b). During freezing, the respiratory rhythm changed in a stereotypic manner and matched the rhythmic head-motion and the prefrontal LFP oscillation (Supplementary Fig. 3c,d). Interestingly, the respiratory peak frequency was distinct for different types of fear behavior and prefrontal peak frequency faithfully matched it, in support to the generality of the described phenomenon (Supplementary Fig. 3e).

To further investigate the extent of respiratory entrainment of prefrontal circuits, we examined the firing of extracellularly recorded single neurons in mPFC in relation to the respiratory phase (**Fig. 2a**). We observed that ~60% of putative prefrontal principal cells (PN) and ~90% of putative inhibitory interneurons (IN), identified based on their extracellular waveform features^28^ (**Supplementary Fig. 4a,b**), were significantly modulated by the phase of respiration cycle (**Fig. 2b-c, Supplementary Fig. 4c**). Most modulated cells fired preferentially in the trough/ascending phase of the local oscillation, corresponding to the inhalation phase (**Fig. 2d, Supplementary Fig. 4d**) and are even more strongly modulated by the phase of the local oscillation (**Fig. 2c, Supplementary Fig. 4e,f**).

**Figure 2.**
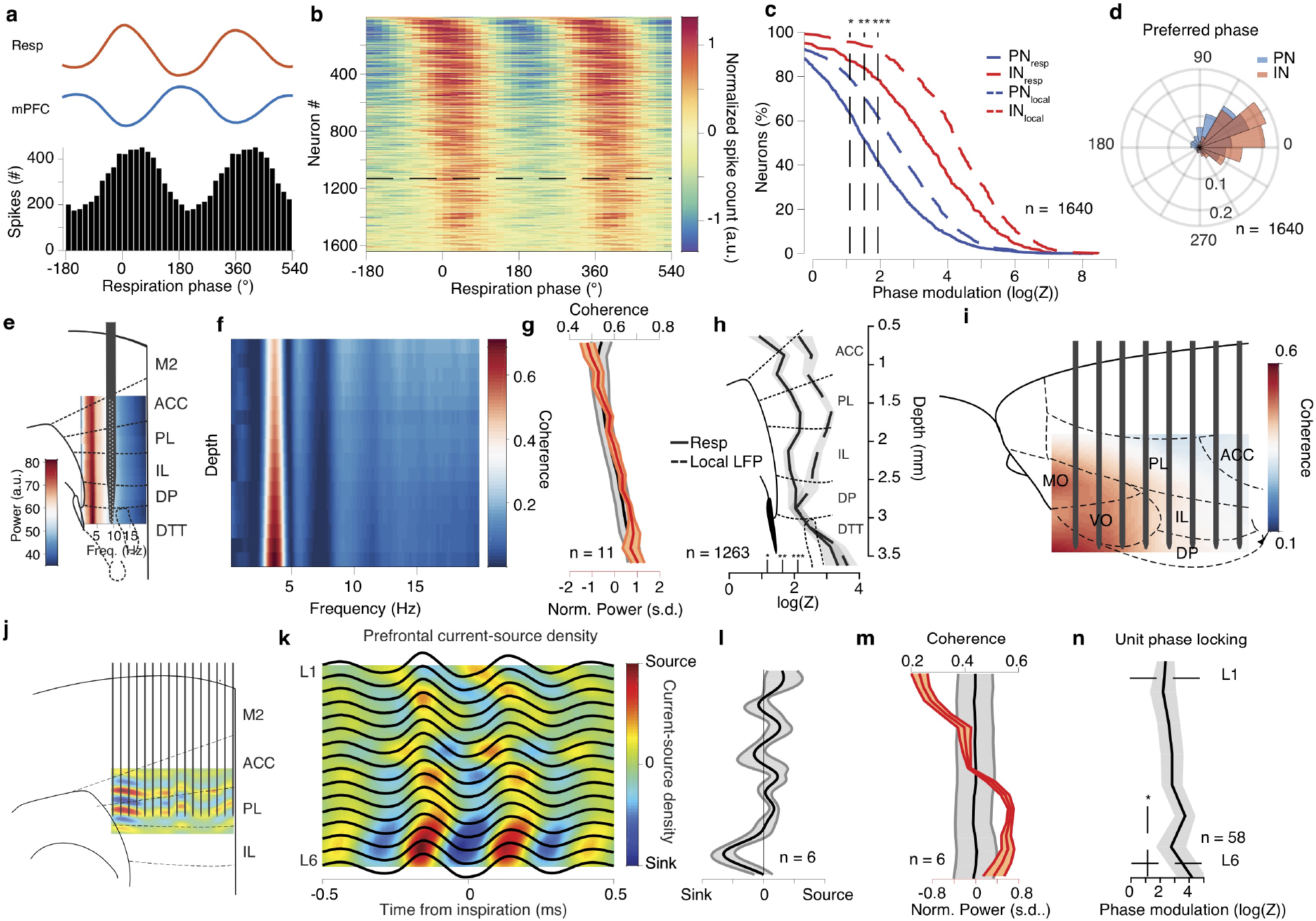
Topography of prefrontal neuronal entrainment by respiration. (**a**) Respiration phase histogram of the spiking activity of an example prefrontal neuron. Top, associated average respiration (red) and mPFC LFP (blue) traces. (**b**) Color-coded normalized phase histograms of all prefrontal neurons, ordered by phase modulation magnitude (n = 1640 neurons, n = 13 mice). The horizontal dashed line indicates the significance threshold for the logZ. (**c**) Cumulative distribution of the logZ for all prefrontal PNs (blue, n = 1250 neurons) and INs (red, n = 390 neurons). Phase modulation is assessed in relation to the respiration (solid lines) and the local prefrontal LFP (dashed lines). (**d**) Distribution of the preferred phase for PNs (blue) and INs (red). The height of each bar corresponds to the relative number of units. (**e**) Schematic depiction of a typical recording using a high-density silicon polytrode inserted in the deep layers of the mPFC, overlaid on an example depth- and frequency-resolved power spectrum spanning all medial prefrontal subregions. (**f**) Example depth- and frequency-resolved coherence between the respiration and local prefrontal LFP spanning all medial prefrontal subregions. (**g**) Average depth-resolved normalized power (red) and coherence in the 2–5 Hz band (black) (n = 11 mice). (**h**) Depth-resolved average phase modulation statistics (logZ) (n = 1263 cells, n = 11 mice). (**i**) Example of 2D coherence between respiration and local LFP throughout the frontal subregions. (**j**) Schematic depiction of a 16-shank probe (50μm shank spacing) inserted in the prelimbic region of the mPFC to record simultaneously from all cortical layers and an example inspiration-triggered current-source density profile. (**k**) Example average inspiration-triggered LFP traces and overlaid corresponding translaminar current-source density profile from the dorsal mPFC. (**l**) Average inspiration-triggered translaminar normalized current-source density profile from the dorsal mPFC (n = 6 mice). (**m**) Average cortical layer-resolved profile of the normalized 2–5 Hz band local LFP power (red) and coherence with respiration (coherence) (n = 6 mice). (**n**) Cortical layer-resolved phase modulation statistics (logZ) (n = 58 cells, n = 6 mice). Shaded areas, mean ± s.e.m. a.u., arbitrary units; s.d., standard deviations; L1, layer 1; L6, layer 6. Stars indicate significance levels (* p<0.05; ** p<0.01; *** p<0.001).

Having established the generality of the coupling between respiration and LFP oscillation in the mPFC across distinct states, consistent with previous reports^21,23,29^, we focused on the enigmatic and least understood quiescence and slow-wave sleep states. To understand the potential role of breathing in orchestrating the hippocampo-cortical dialogue and supporting memory consolidation, we undertook a detailed investigation of the mechanism and function of the respiratory modulation.

### Topography of respiratory entrainment

mPFC consists of multiple subregions along the dorsoventral axis, all of which are characterized by a differential afferent and efferent connectivity and behavioral correlates^30,31^. To understand the origin and anatomical substrate of the respiratory entrainment of prefrontal circuits, we performed a three-dimensional trans-laminar and trans-regional characterization of the mPFC field potentials using custom-designed high-density silicon probes in freely-behaving and head-fixed mice (**Fig. 2e-n**). These recordings revealed a consistent increase from dorsal to ventral mPFC in both the power of the respiration-related oscillation and its coherence with the respiration (**Fig. 2e-g**). Similarly, the coherence was stronger in the anterior prefrontal regions (**Fig. 2i**).

The presence of an oscillation in the mPFC with this particular profile could also be consistent with a volume conducted signal from the high amplitude field potentials generated by bulbar dipoles, since olfactory bulb (OB) LFP is dominated by respiration-related oscillations^32–34^. To examine this hypothesis, we recorded LFP activity across the prefrontal cortical layers and calculated the current-source density (CSD) (**Fig. 2j-l, Supplementary Fig. 4g**). This analysis revealed a prominent pattern of sinks in the deep cortical layers at the inspiration phase giving rise to an increased LFP power and unit-LFP coupling in the deep layers (**Fig. 2k-n**), weighing against the hypothesis of volume conduction and suggestive of a synaptic origin of the prefrontal LFP oscillation.

Harnessing the advantages of spatial information from the silicon probe recordings, we characterized the entrainment of single units across prefrontal subregions and cortical layers. Although cells were phase-modulated throughout the prefrontal subregions, the average modulation strength was increased as a function of distance from the dorsal surface (**Fig.2h, Supplementary Fig. 4e,f**). These results, given the increased density of polysynaptic projections from the OB to ventral mPFC subregions^23^, suggest that the bulbar reafferent input to the mPFC is giving rise to the observed LFP signals. Rhythmic air flow could entrain the olfactory sensory neurons (OSNs), that are known to respond both to odors and mechanical stimuli^35^, and propagate through the olfactory bulb and the olfactory system to the prefrontal region.

### Widespread respiratory modulation of limbic circuits

Given the prominent modulation of mPFC by respiration, we hypothesized that a concurrent entrainment of other limbic regions could be underlying their generalized long-range interaction, as has been suggested before for theta oscillations^6,36^. For this, we turned our attention to other limbic structures, reciprocally connected to the mPFC that are known to interact with prefrontal networks in different behaviors. Using large-scale single-unit and laminar LFP recordings from the dorsal hippocampus, we identified that in both dorsal CA1 and dentate gyrus (DG), ~60% of PNs and 80% CA1 INs were modulated by the phase of respiration, firing preferentially after the inspiration (**Fig. 3a-c**), in line with previous reports of respiratory entrainment of hippocampal activity^21,22,37,38^. A separation of CA1 PNs based on their relative position within the pyramidal layer into populations with known distinct connectivity patterns^39,40^ did not reveal particular differences in their modulation by respiration, suggesting a generality of this entrainment throughout the CA1 sub-populations (**Supplementary Fig. 5b**).

**Figure 3.**
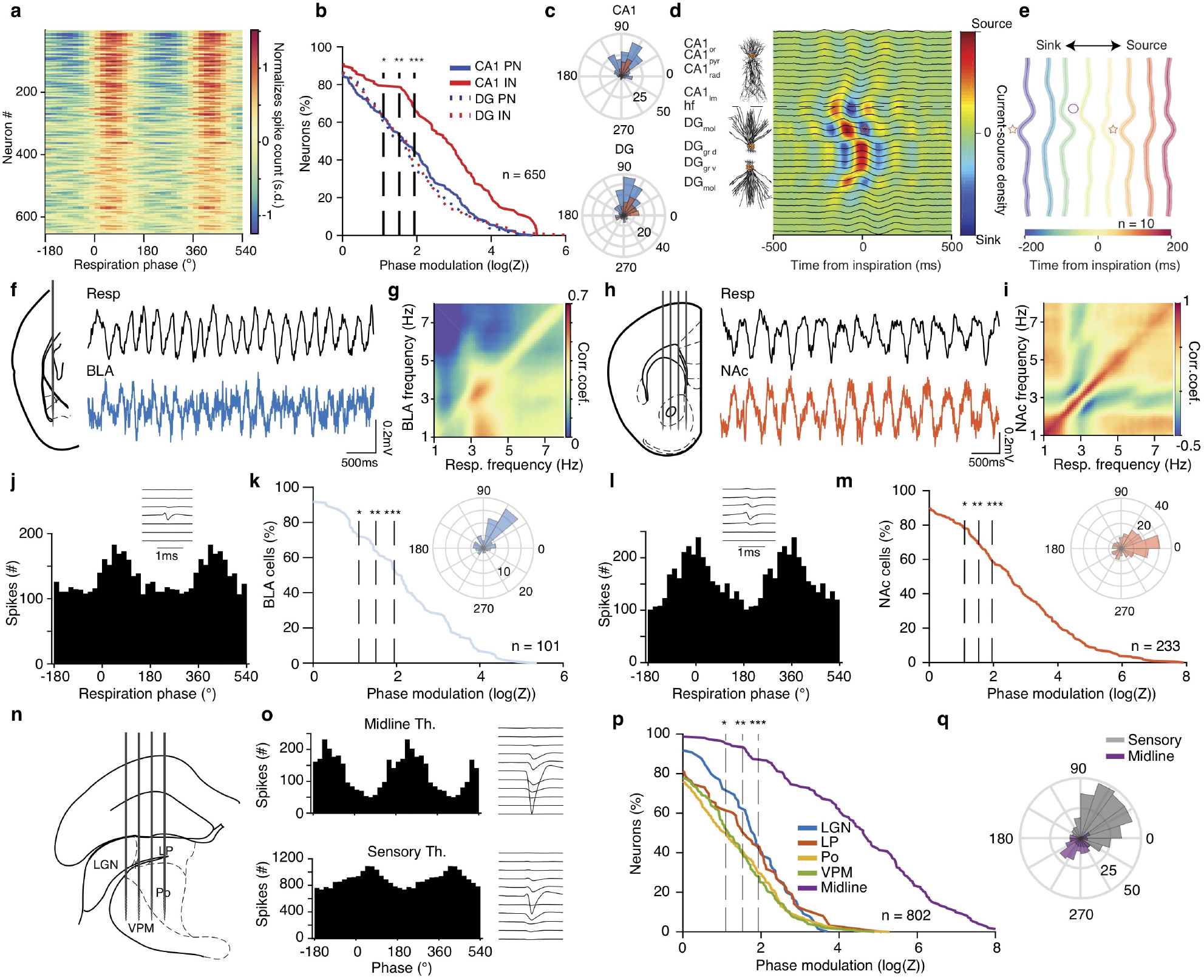
Breathing modulates BLA, NAc, hippocampal and thalamic neuronal activity. (**a**) Color-coded respiration phase histograms of the normalized firing rate of all hippocampal cells (n = 650 cells). (**b**) Cumulative distribution of the modulation strength for all CA1 and DG PNs (CA1, n = 226 cells; DG, n = 206 cells) and INs (CA1, n = 98 cells; DG, n = 120 cells). (**c**) Distribution of the preferred phase for all significantly phase locked CA1 (top) and DG (bottom) cells. (**d**) Schematic depiction of CA1 pyramidal and DG granular cell somato-dendritic domains aligned to the example inspiration-triggered high-density CSD profile of the dorsal hippocampus. Horizontal dashed line indicates the hippocampal fissure. (**e**) Average normalized inspiration-triggered CSD profile of dorsal hippocampus at different lags from inspiration (n = 10 mice). Stars mark the middle molecular layer sink and circles the outer molecular layer sink (**f**, **h**) Left, schematic of recording configurations. Right, example simultaneously recorded respiration and BLA (**f**) or NAc (**h**) LFP trace. (**g**, **i**) Example frequency-resolved comodulation of respiration and BLA (**g**) or NAc (**i**) LFP oscillation power. (**j**, **l**) Respiration phase histogram of the spiking activity of one example BLA (**j**) or NAc (**l**) neuron. Insets, the spatio-temporal spike waveform for the respective units. (**k**, **m**) Cumulative distribution of the logZ for all BLA (**k**) or NAc (**m**) cells (BLA: n = 101 cells, NAc: n = 233 cells). Insets, distribution of the mean preferred respiration phases of all significantly modulated cells. (**n**) Schematic of recording configuration for sensory thalamus. (**o**) Respiration phase histograms of the spiking activity (left) and spatio-temporal spike waveforms of respective example units from the midline (top) and sensory (bottom) thalamus. (**p**) Cumulative distribution of the modulation strength for all thalamic neurons (n=802 cells). (**q**) As in (**e**) but for sensory and midline thalamus. Stars indicate significant phase modulation levels (* P<0.05; ** P<0.01; *** P<0.001).

To understand what afferent pathways are responsible for breathing-related synaptic currents that underlie the modulation of spiking activity, we calculated finely-resolved (23μm resolution) laminar profile of inspiration-triggered dorsal hippocampal current-source density, enabling the identification of synaptic inputs into dendritic sub-compartments. Although the LFP profile only highlights the prominence of the respiratory band in the DG hilus region (Supplementary Fig. 5a), high-resolution CSD analysis revealed the presence of two distinct and time-shifted respiratory-related inputs in DG dendritic sub-compartments (**Fig. 3d, Supplementary Fig. 5d**). Inspiration was associated with an early sink in the outer molecular layer of DG, indicative of an input from the layer II (LII) of the lateral entorhinal cortex (LEC), followed by a sink in the middle molecular layer of DG, indicative of an input from the layer II of medial entorhinal cortex (MEC) (**Fig. 3d,e**).

To explore the extent of limbic entrainment by respiration, we further recorded LFP and single-unit activity in the BLA, NAc as well as somatic and midline thalamus (**Fig. 3f-q**). Similar to mPFC, LFP in both BLA and NAc was comodulated with breathing across a range of frequencies, with most prominent modulation at ~4 Hz, the main mode of breathing frequency during quiescence (**Fig. 3f-i**), and exhibited reliable cycle-to-cycle phase relationship with the respiratory oscillation (**Supplementary Fig. 6c,e**). Given the nuclear nature and lack of lamination of these structures, which obfuscates the interpretation of slow LFP oscillations, we examined the modulation of single-units by the phase of breathing. Phase-modulation analyses of the spiking activity revealed that a large proportion of BLA, NAc, and thalamic neurons are modulated by respiration, firing in distinct phases of the breathing cycle (**Fig. 3j-q**).

### Reafferent origin of limbic gamma oscillations

A prominent feature of prefrontal cortex LFP are fast gamma oscillations (~80 Hz)^41,42^ (**Fig. 4a**). To investigate the relationship of prefrontal gamma oscillations to the breathing rhythm and well-known OB gamma oscillations of similar frequency^34,43–45^ (Supplementary Fig. 6a), we recorded simultaneously from the two structures and calculated the phase-amplitude coupling between breathing and gamma oscillations (**Fig. 4a,b, Supplementary Fig. 6a**). Both OB and mPFC fast gamma oscillations are modulated by respiratory phase and gamma bursts occur predominantly simultaneously and in the descending phase of the local LFP (**Fig. 4c**). The simultaneously occurring OB and mPFC gamma oscillations match in frequency and exhibit reliable phase relationship with a phase lag suggesting directionality from the OB to the mPFC (**Fig. 4d, Supplementary Fig. 6b**). To examine the underlying synaptic inputs mediating the occurrence of these oscillations in the mPFC, we calculated CSD across mPFC layers, triggered on the phase of the OB gamma bursts. This analysis revealed a discrete set of sinks across prefrontal layers associated with OB bursts (**Fig. 4i,j**). Similar to the slow time scale LFP signals, these results suggest that fast gamma oscillations in mPFC are generated by OB-gamma rhythmic polysynaptic inputs to mPFC and are not a locally generated rhythm.

**Figure 4.**
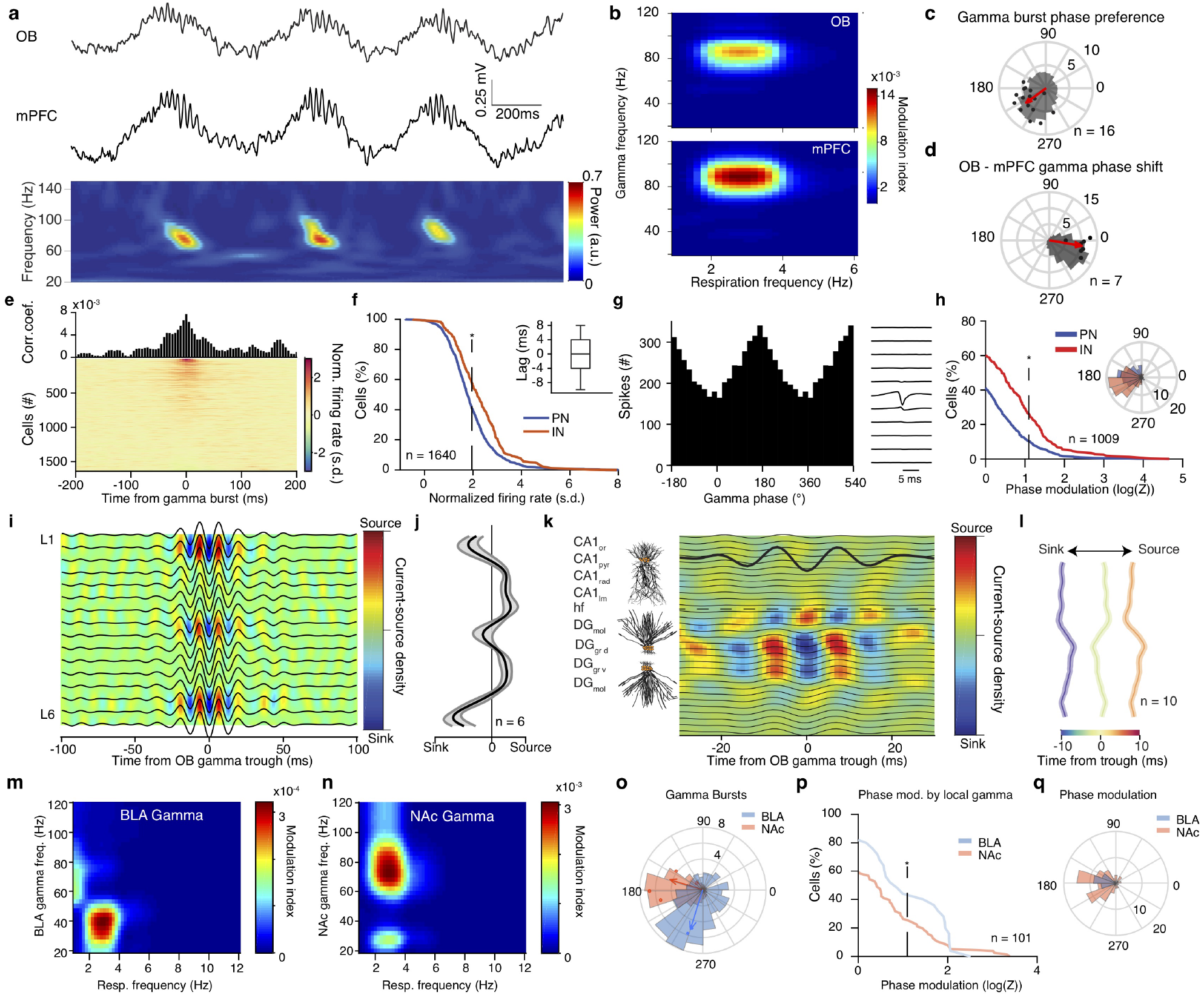
Reafferent gamma entrainment of limbic circuits. (**a**) Example simultaneously recorded LFP traces (top) from OB and mPFC LFP and its spectral decomposition in the gamma range (bottom). (**b**) Color-coded modulation strength of OB (top) and mPFC (bottom) gamma power by respiration phase for an example animal. (**c**) Phase distribution of mPFC gamma bursts for an example animal (gray histogram) and average preferred phase and phase modulation strength (logZ) for all animals (n = 16 mice). The red arrow indicates the population average preferred phase and log(Z). (**d**) Distribution of the phase shift between OB and mPFC gamma filtered traces for one example animal (gray histogram) and average phase shift and phase-coupling strength (log(Z), red dots) for all animals (n = 7 mice). (**e**) Gamma-burst triggered time histogram for one example mPFC cell and color-coded normalized time histograms for all mPFC cells (n = 1640 cells). (f) Cumulative distribution of the gamma-triggered normalized firing of mPFC PNs (n = 1250 cells) and INs (n = 390 cells). Inset, time lag between time from gamma burst and peak firing probability for all significantly responsive cells. (**g**) Gamma phase histogram of one example mPFC unit (left) and the respective unit spike spatio-temporal waveform (right). (**h**) Cumulative distribution of the modulation strength (logZ) for all PNs (blue, n = 685 neurons) and INs (red, n = 324 neurons). Phase modulation is assessed in relation to the phase of the locally recorded prefrontal gamma oscillation. Inset, distribution of the mean preferred phases of all significantly modulated PN and IN cells. (**i**, **j**) Example (**i**) and average zero-lag (**j**) OB gamma-triggered translaminar CSD of the dorsal mPFC LFP profile. (n = 6 mice). (**k**) Example OB gamma-triggered CSD profile of dorsal hippocampus. Horizontal dashed line indicates the hippocampal fissure. (**l**) Average normalized OB-triggered current-source density profile of dorsal hippocampus at different time lags from OB gamma trought (n = 10 mice). (**m**, **n**) Example phase-power modulation of BLA (**m**) and NAc (**n**) gamma activity by respiration. (**o**) Example distribution of the respiratory phase of BLA and NAc gamma bursts (histogram) and mean preferred phase of gamma occurrence and modulation strength (dots; BLA, blue, n = 3 mice; NAc, red, n = 4 mice). (**p**) Cumulative distribution of modulation strength for local gamma phase entrainment of spikes of all BLA (blue, n = 25 cells) and NAc cells (red, n = 76 cells). (**q**) Distribution of the mean preferred gamma phase for each significantly modulated BLA and NAc cell. Star indicates significance (* P<0.05). Shaded areas, mean ± s.e.m.

Examining the OB ~80 Hz gamma-triggered dorsal hippocampal CSD reveals a DG outer molecular layer sink, indicative of an OB gamma propagating to DG via the LEC LII input (**Fig. 4k,l**), a profile distinct from the similar frequency CA1lm gamma (Supplementary Fig. 5c). In parallel, slow BLA gamma (~40 Hz) and fast NAc gamma (~80 Hz) oscillations are modulated by the phase of breathing, occurring predominantly in the trough and ascending phase of breathing respectively (**Fig. 4m-o**).

To examine whether these respiration-modulated OB-mediated gamma oscillations have a functional role in driving local neuronal activity, we quantified coupling of local single units to mPFC gamma signals, revealing that ~40% of principal cells and ~55% of interneurons increased their firing rate in response to local gamma oscillations (**Fig. 4e,f**). Interestingly, ~10% of PN and ~30% of IN were significantly phase modulated by gamma oscillations, firing preferentially in the trough of the local oscillation (**Fig. 4g,h**). Similarly, ~40% of BLA and ~25% of NAc cells fire preferentially in the trough of the local gamma oscillations (**Fig. 4p, Supplementary Fig. 6d,f**). Thus OB gamma propagates

### Efferent and reafferent mechanisms of respiratory entrainment

These results suggest a mechanistic picture in which the OB reafferent gamma and respiration-locked currents are responsible for the observed respiration-associated LFP patterns in the mPFC, consistent with disruption of these LFP patterns after OB lesion or tracheotomy^20,23,29,46^. However, the distributed and massive modulatory effect that respiration had on unit activity in these regions is at odds with the anatomically-specific synaptic pathways that we identified as responsible for slow and fast currents.

To causally test whether OB reafferent input is the sole origin of the LFP patterns and unit entrainment, we resorted to a pharmacological approach, that enables selective removal of the reafferent input. A well-characterized effect of systemic methimazole injection is the ablation of the olfactory epithelium that hosts the olfactory sensory neurons^47^, known to express mechanoreceptors^35^. Effectively, this deprives the OB of olfactory and respiratory input, while leaving the bulbar circuits intact, enabling us to study the activity of the de-afferentiated brain in freely-behaving mice. This manipulation eliminated the respiration-coherent prefrontal oscillatory LFP component (**Fig. 5a-d, Supplementary Fig. 7a-c**), consistent with the disappearance of the CSD sink in deep layers (**Fig. 5e**), while at the same time abolishing the olfactory-related prefrontal gamma oscillations (**Fig. 5f,g**). These results confirm the hypothesis that a respiratory olfactory reafference (ROR) is responsible for the rhythmic cortical LFP, as suggested previously^20,23,29,46^.

**Figure 5.**
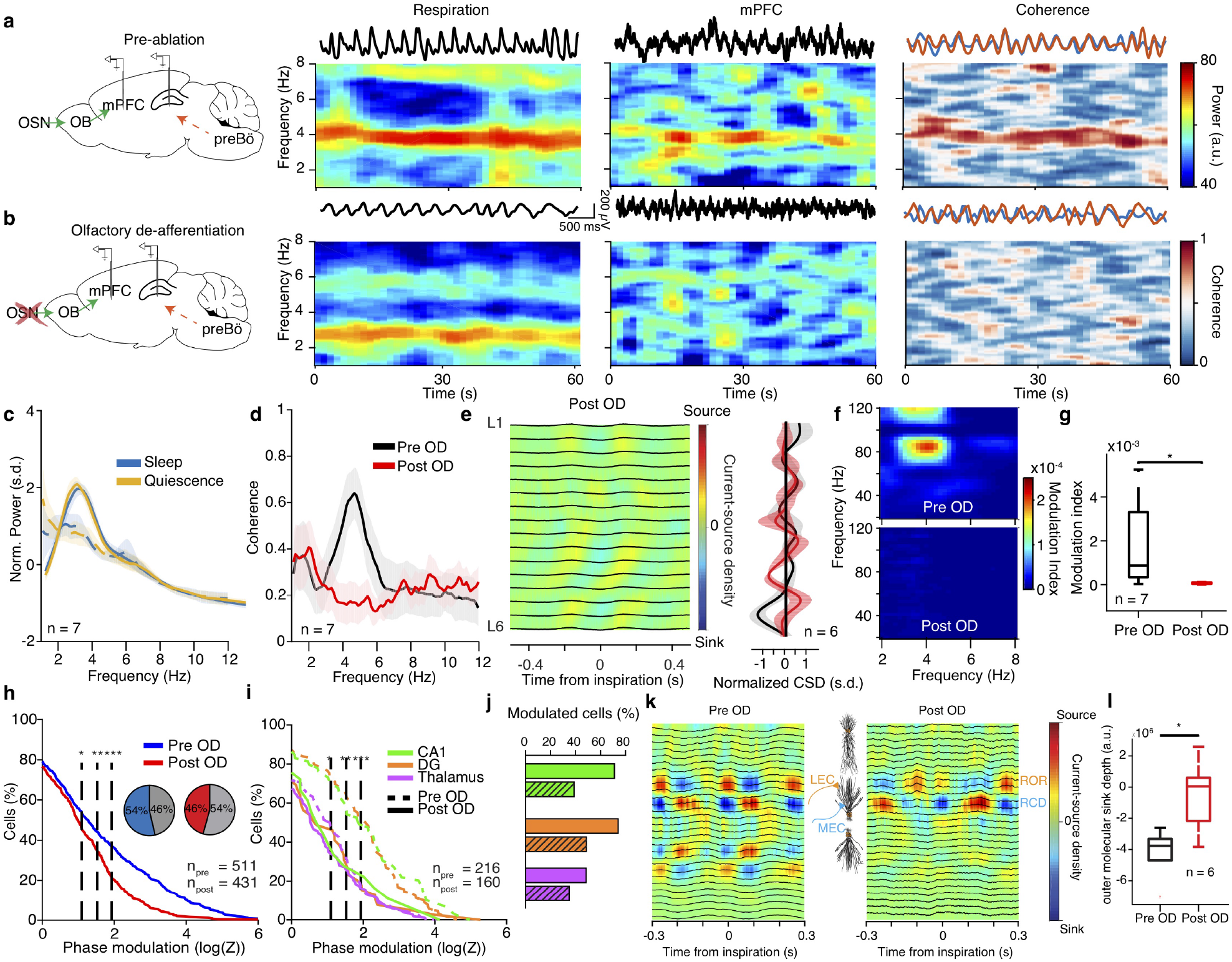
Reafferent respiratory input accounts for LFP but not neuronal entrainment. (a,b) Left, schematic of the manipulation strategy. Right, example time-frequency decomposition of power and coherence between respiratory and mPFC LFP signals during quiescence before (**a**) and after (**b**) OD. (**c**) Average normalized power spectra before and after OD (n = 7). (**d**) Coherence spectrum between respiration and mPFC LFP before and after OD (n = 7 mice). (**e**) Left, example inspiration-triggered CSD of the mPFC LFP during quiescence and sleep after OD. Right, average normalized CSD at zero lag (n = 6 mice). (f) Example power-phase modulation of mPFC gamma oscillations before (top) and after (bottom) OD. (**g**) Average mPFC power-phase modulation strength of ~80 Hz gamma oscillations (n = 7 mice; paired t-test: before vs. after OD). (**h**) Cumulative distribution of modulation strength for all mPFC neurons pre and post OD (Pre: n = 511 cells; Post: n = 431 cells). Inset, percentage of significantly phase-modulated cells before and after OD. (**i**) Cumulative distribution of modulation strength for CA1, DG and somatic thalamus neurons before and after OD. (**j**) Percentage of significantly phase-modulated cells before and after OD. (**k**) Example inspiration-triggered CSD of the dorsal hippocampus LFP before (left) and after (right) OD. (**l**) Average outer molecular layer sink depth (n = 6 mice; paired t-test: before vs. after OD). Shaded areas, mean ± s.e.m. a.u., arbitrary units; s.d., standard deviations; n.s., not significant. Shaded areas, mean ± s.e.m. Stars indicate significance levels (* P<0.05; ** P<0.01; *** P<0.001). s.d., standard deviations; a.u., arbitrary units; OD, olfactory de-afferentiation.

Surprisingly however, the olfactory de-afferentiation (OD) left most prefrontal and thalamic neurons modulated by breathing, although the strength of modulation was somewhat reduced (**Fig. 5h,i**), suggesting that a so-far undescribed ascending respiratory corollary discharge (RCD) signal, likely propagating from the brainstem respiratory rhythm generators, is responsible for the massive entrainment of prefrontal neurons. Interestingly, following OD, mice exhibited intact memory and fear expression, suggesting that the RCD might be underlying the behavioral expression (Supplementary Fig. 7d-f). Dorsal hippocampal neurons were somewhat stronger affected by the ablation, yet >40% of cells were still significantly phase locked (**Fig. 5i,j**), indicating a differential degree of contribution of RCD and ROR to unit firing across mPFC, HPC and thalamus. In contrast to the prefrontal CSD, the olfactory de-afferentiation led to a strong reduction of the outer molecular layer current-sink originating in LEC LII in the respiration-locked CSD (**Fig. 5k,l**), while leaving MEC LII sink and other non-respiration related CSD patterns intact (Supplementary Fig. 8c), suggesting that the LEC input is driven by ROR, while MEC input is driven by RCD.

### Hippocampal network dynamics are modulated by breathing

From the results so far, it is clear that hippocampal neuronal activity is massively modulated by breathing, by means of entorhinal inputs to the DG. However, during quiescence and slow-wave sleep, hippocampal activity is characterized by recurring nonlinear population events such as dentate spikes (DS) and sharp-wave ripple complexes.

CA1 ripples are local fast oscillations in the pyramidal layer of CA1, generated by the rhythmic interplay between PNs and INs in response to strong depolarization from CA3 population spikes, identified as sharp negative potentials (sharp-waves) in the CA1 stratum radiatum^48^ (**Fig. 6a-c, Supplementary Fig. 8a**). During ripples, CA1 PN and IN, and to a lesser extent DG cells, were strongly activated (**Fig. 6d**). Ripple occurrence was strongly modulated by breathing biased towards the post-inspiratory phase, while ripples were suppressed during exhalation (**Fig. 6e-g**). Following olfactory de-afferentiation, ripples remained sig-nificantly phase modulated by breathing (**Fig. 6h**), suggesting that RCD is the main source of their modulation. Keeping up with the role of entorhinal input mediating respiratory drive on ripples, we observed a consistent relationship between the mag-nitude of MEC LII sink directly preceding ripple occurrence and the phase within the respiratory cycle of the ripple occurrence (**Supplementary Fig. 8c,d**).

**Figure 6.**
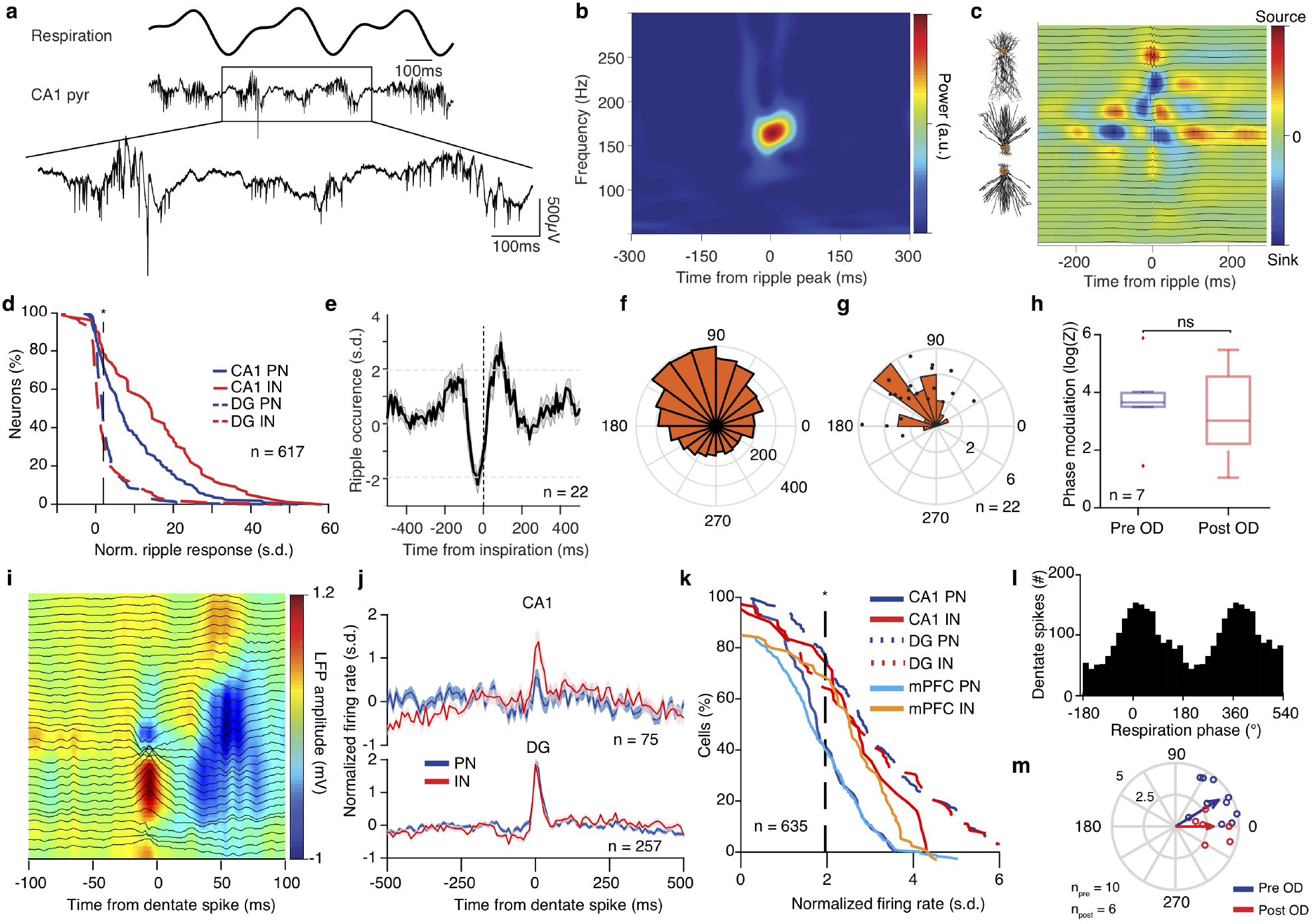
Breathing modulates hippocampal ripples and dentate spikes. (**a**) Example traces of the respiratory signal and CA1 pyramidal layer LFP. In the magnified LFP signal, ripple events and the associated spiking activity can be observed. (**b**) Average ripple-triggered time-frequency wavelet spectrogram of the CA1 pyramidal layer LFP from one example animal. (**c**) Schematic of the CA1 pyramidal and granular cells somato-dendritic domains aligned to the average ripple-triggered CSD profile of the hippocampal LFP activity for one example animal. (**d**) Cumulative distribution of the ripple-triggered normalized firing of CA1 and DG PNs (CA1, n = 220 cells; DG, n = 202 cells) and INs (CA1, n = 76 cells; DG, n = 119 cells). (**e**) Average cross-correlation between inspiratory events and ripple occurrence (n = 22 mice). Dashed horizontal lines indicate the significance levels. (f) Distribution of the respiratory phase of occurrence of individual ripple events for one example animal (n = 4813 ripples). (**g**) Distribution of average phase and modulation strength for ripples (n = 22 mice). (**h**) Phase modulation of ripples before and after OD. (**i**) Color-coded LFP power depth profile and overlaid LFP traces of an example dentate spike. (**j**) Average dentate spike triggered normalized spiking activity for all CA1 (top) and DG (bottom) PNs and INs. (**k**) Cumulative distribution of the dentate spike-triggered normalized firing of CA1, DG and mPFC PNs (CA1, n = 44 cells; DG, n = 175 cells; mPFC, n = 239 cells) and INs (CA1, n = 31 cells; DG, n = 82 cells; mPFC, n = 64 cells). (**l**) Respiration phase histogram for dentate spikes in one example animal. (**m**) Preferred respiratory phase of dentate spike occurrence and phase modulation strength for all animals (dots) and population average (arrows) before and after OD (*n_pre_* = 10 mice,*n_post_* = 6 mice). s.d., standard deviations; a.u., arbitrary units; n.s., not significant; OD, olfactory de-afferentiation. Shaded areas, mean ± s.e.m. Stars indicate significance levels (* P<0.05; ** P<0.01; *** P<0.001).

Another prominent hippocampal offline state-associated pattern are dentate spikes, defined as large positive potentials in the DG hilus region^49^ which are believed to occur in response to strong inputs in the molecular layer, such as during entorhinal UP states (**Fig. 6i, Supplementary Fig. 5d**). During DS, both DG (~70% PN and IN), CA1 (~40% PN and ~70% IN) and mPFC (~40% PN and ~70% IN) cells were strongly excited (**Fig. 6j,k**). Consistent with respiratory entrainment of the entorhinal inputs, the occurrence of DS was strongly modulated by the breathing phase both before and after OD, with the majority of events occurring after inspiration (**Fig. 8l,m**).

### Breathing organizes prefrontal UP states and hippocampal output

Similar to the hippocampus, during quiescence and slow-wave sleep neocortical circuits exhibit nonlinear bistable dynamics in the form of DOWN and UP states. We posited that the strong rate modulation of prefrontal neural activity by breathing would bias these dynamics. To test this prediction, we identified prefrontal UP and DOWN states based on the large-scale population activity and characterized their relationship with the phase of the ongoing breathing rhythm (**Fig. 7a**). Surprisingly, both UP and DOWN state onsets were strongly modulated by the breathing phase and time from inspiration (**Fig. 7b-d**), while the magnitude of UP states followed the profile of UP state onset probability (**Fig. 7e**). In line with the results on ripples and prefrontal units, UP and DOWN state modulation was not affected by olfactory de-afferentiation, suggesting that RCD is the source of this modulation (**Fig. 7f,g**).

**Figure 7.**
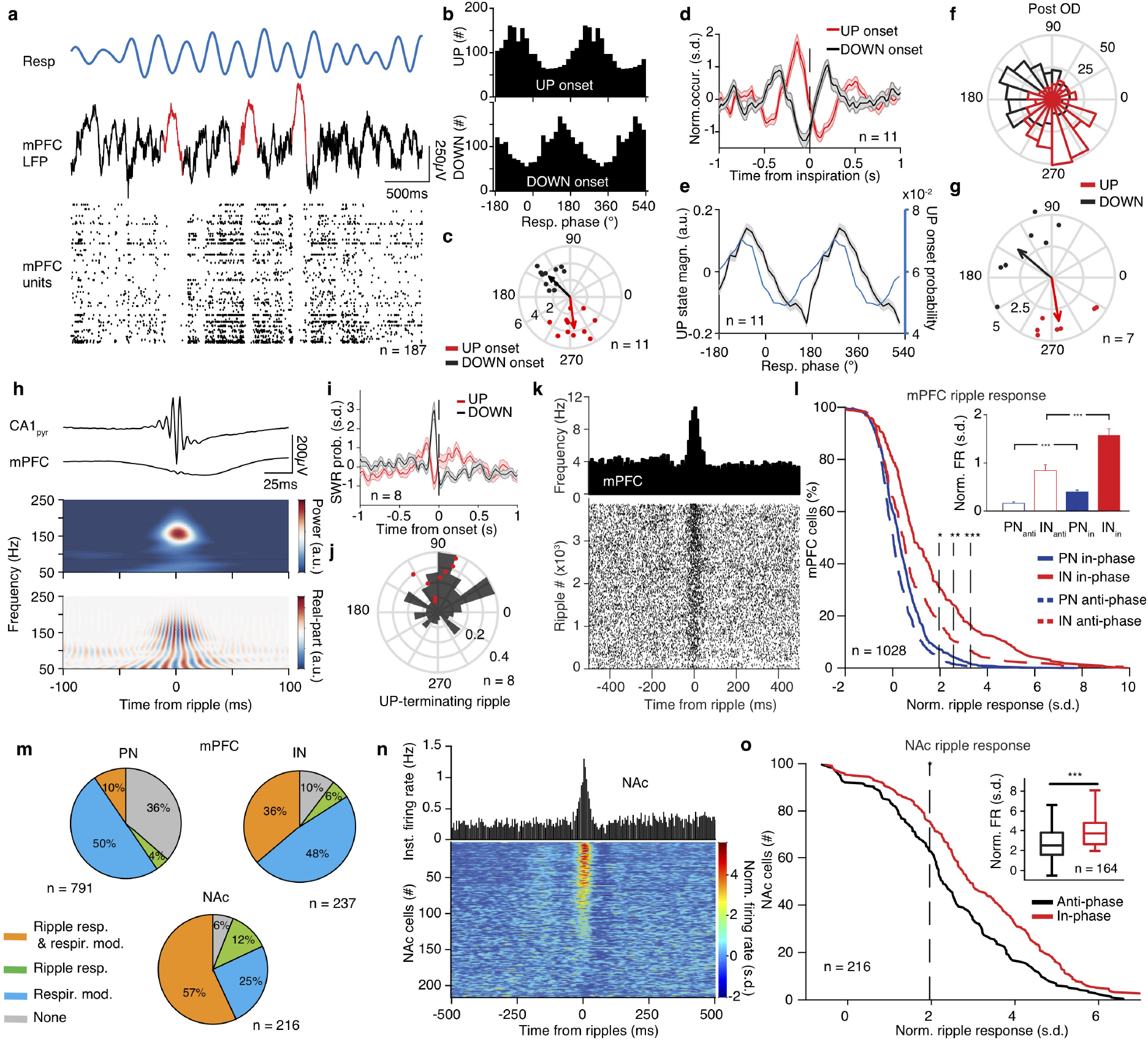
Breathing organizes prefrontal UP states and modulates hippocampal output. (**a**) Example traces of respiration signal (top), mPFC LFP (middle trace) and spike trains of 183 simultaneously recorded mPFC units during sleep. Three typical delta waves and the corresponding DOWN states of the neuronal population are marked with red. (**b**) Example distributions of the breathing phase of UP (top) and DOWN (bottom) state onsets. (**c**) Distribution of preferred breathing phase of UP and DOWN states (n = 11 mice). (**d**) Cross-correlation of UP and DOWN state onsets with respect to inspiration. (**e**) Normalized power (black) and occurrence probability (blue) for prefrontal UP states (n = 11 mice). (**f**, **g**) Example (**f**) and population (**g**) distribution of preferred breathing phase of UP and DOWN state occurrence after OD (n = 7). (**h**) Example average ripple-triggered CA1pyr and mPFC LFP traces and wavelet spectral decomposition of mPFC LFP power (upper) and real-part (lower) (n = 3162 ripples). (**i**) Probability of SWR occurrence as a function of time from UP or DOWN state onset (n = 8 mice). (**j**) Example (black) and distribution of preferred breathing phase of SWR occurrence, for SWRs that are terminating an UP state (red dots, n = 8 mice). (**k**) Example mPFC unit spiking raster across individual ripples (bottom) and cross-correlogram of unit firing to ripple (top). (**l**) Cumulative distribution of the ripple-triggered normalized firing of mPFC PNs (n = 791 cells) and INs (n = 237 cells) in response to ripples occurring in the preferred (in-phase) and in the least-preferred (anti-phase) phases of respiration (n = 11 mice). Inset, average ripple-triggered firing rate (PN: 791 cells. IN: 237 cells) (Wilcoxon signed-rank test, in-phase versus anti-phase). (**m**) Top, pie charts indicating the percentage of all mPFC PNs (left; n = 791 cells) and INs (right; n = 237 cells) that are either phase modulated by respiration (resp. mod), responding significantly to ripples (ripple resp.), being both significantly modulated by breathing and significantly responsive to ripples or neither. Bottom, similarly, for all NAc cells (n = 216 cells). (**n**) Cross-correlation of firing with respect to ripple time for one example NAc unit (top) and color-coded cross-correlograms for all NAc cells (n = 216 cells). (**o**) Cumulative distribution of the ripple-triggered normalized firing of NAc cells (n = 216 cells) in response to, as in (**l**), “in-phase” and “anti-phase” ripples (n = 4 mice). Inset, average ripple-triggered firing rate (n = 164 cells, Wilcoxon signed-rank test, in-phase versus anti-phase). Shaded areas, mean ± s.e.m., s.d., standard deviations; a.u., arbitrary units; OD, olfactory de-afferentiation; FR, firing rate. Stars indicate significance levels (* P<0.05; ** P<0.01; *** P<0.001).

Previous observations during sleep in rats identified a correlation between ripple occurrence and cortical SO^11,50,51^. Here, ripples preceded the termination of prefrontal UP states and onset of the DOWN states both before (**Fig. 7i**) and after de-afferentation (**Supplementary Fig. 8e**), with ripples contributing to UP state termination occurring in the early post-inspiratory phase (**Fig. 7j**). This is in line with the RCD-driven synaptic inputs to the DG middle molecular layer preceding ripple events, which are suggesting an RCD-mediated coordination of SWR occurrence with the MEC UP states (**Supplementary Fig. 8c,d**).

Ripple output is known to recruit prefrontal neural activity^50,51^. In line with this, hippocampal ripples evoked a response in prefrontal LFP and gave rise to an efferent copy detected as a local increase in fast oscillatory power in the PFC LFP^52^ (**Fig. 7h**). In response to ripple events, ~14% of prefrontal PNs and ~42% of INs exhibited increased firing (**Fig. 7k,l, Supplementary Fig. 8f**). In parallel, ~69% of NAc cells are significantly driven by ripple events (**Fig. 7n,o, Supplementary Fig. 8f**). Interestingly, in both mPFC and NAc there is a great overlap between cells that are phase modulated by breathing and that are responsive to ripples (**Fig. 7m**), while ripples occurring in the preferred post-inspiratory phase of ripple occurrence, drive a greater fraction of cells from both structures to fire significantly more compared to anti-preferred phase (**Fig. 7l,o**).

## Discussion

In this study, we demonstrate that the respiratory rhythm provides a unifying global temporal coordination of neuronal firing and nonlinear dynamics across cortical and subcortical limbic networks. Using recordings of the three-dimensional LFP profile of the mPFC (**Fig. 1,2**) and large-scale unit recordings from the hippocampus, amygdala, nucleus accumbens and thalamus (**Fig. 3**), we identified that during quiescence and slow-wave sleep limbic LFPs are dominated by breathing-related oscillatory activity, while the majority of neurons in each structure are modulated by the phase of the respiratory rhythm and reafferent gamma oscillations (**Fig. 4**). Using pharmacological manipulations paired with large-scale recordings, we causally identified a joint mechanism of respiratory entrainment (**Fig. 8a**), consisting of an efference copy of the brainstem respiratory rhythm (respiratory corollary discharge; RCD) that underlies the neuronal modulation and a respiratory olfactory reafference (ROR) that contributes to the modulation and accounts for LFP phenomena (**Fig. 5**). Hippocampal SWR and dentate spikes (**Fig. 6**), as well as prefrontal UP and DOWN states (**Fig. 7**), were strongly modulated by the respiratory phase with RCD being a sufficient source of modulation. This modulation accounts partially for the coordination of hippocampo-cortical nonlinear dynamics (**Fig. 7**). Finally, we comprehensively characterized the distinct phase relationship between different network events within the breathing cycle across limbic structures, painting the picture of how this organization enables the multiplexing and segregation of information flow across the limbic system and providing the basis for mechanistic theories of memory consolidation processes enabled by the respiratory modulation (**Fig. 8b**).

**Figure 8.**
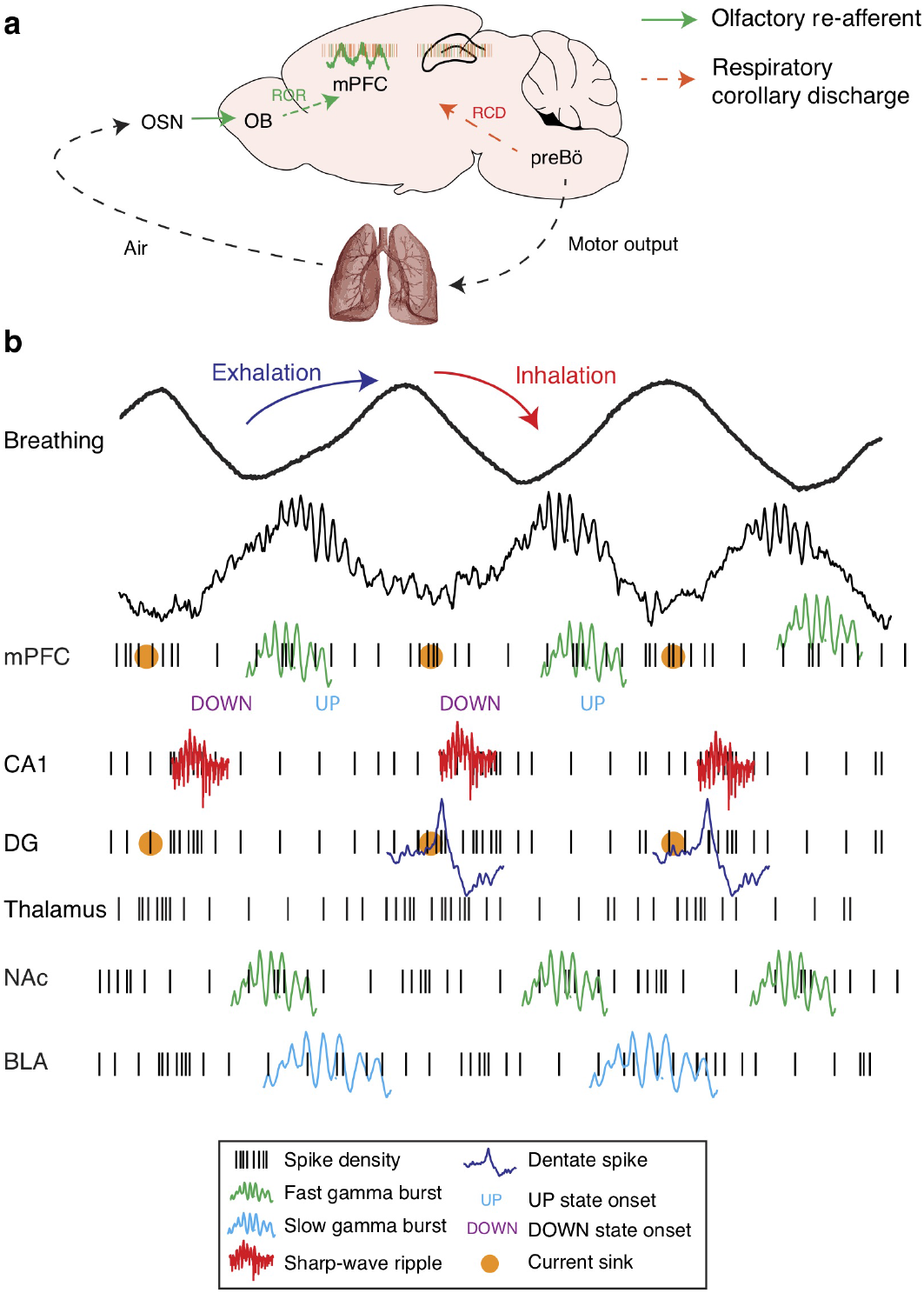
Breathing organizes network dynamics across limbic structures. (**a**) Schematic depiction of the efferent copy pathway carrying the respiratory corollary discharge (RCD) signal and the reafferent pathway carrying the respiratory olfactory reafferent (ROR) signal. (**b**) Summary schematic of the network dynamics organized by breathing throughout all structures studied. Black traces: LFPs; Black ticks: neuronal spikes; Green traces: Fast (~80Hz) gamma; Cyan traces: Slow gamma (~40l·lz); Red traces: CA1 Ripples; Blue traces: Dentate spikes; Orange dots: CSD sinks (mPFC deep layers and DG middle molecular layer).

Over the years, mounting evidence have suggested that breathing can entrain frog^53^ and human EEG^54,55^, as well as LFP and spiking activity in the hedgehog OB and cortex^32^, rodent OB^33–35,56^, cortex^20,23,29,57–59^ and the hippocampus in cats^60^, rodents^21,22,37,38,61^, and humans^24,25^. With this study, we contribute to the ongoing effort to understand the mechanism and role of the respiratory entrainment of brain circuits.

Our results extend and provide a mechanistic explanation and interpretation of the previous studies that described respiration-related LFP oscillations in different brain regions. Here, we leverage large-scale recordings from multiple limbic regions and thousands of cells to comprehensively characterize and uncover the underlying mechanisms of the limbic respiratory entrainment and understand the role of this phenomenon in organizing neuronal activity and coordinating network dynamics between remote regions.

In this study, we expanded the characterization of respiratory entrainment to the NAc, BLA, and thalamus for the first time, while we comprehensively characterized the neuronal entrainment across prefrontal layers and subregions. These results highlight the extent and significance of this modulation, given the crucial role of the interaction between these structures for emotional processing and regulation. We further report the reafferent OB origin of local gamma dynamics and their modulation by breathing, as well as the relation between local gamma oscillations and neuronal activity in all structures. This sheds new light onto the origin and role of prefrontal^41,42,59^, BLA^62^, and NAc^63^ gamma oscillations, that might provide a temporally-optimized privileged route for olfactory reafferent input to affect the ongoing activity, in line with recent reports in humans^24^.

Using a pharmacological olfactory de-afferentiation approach, paired with large-scale recordings, we identified that although cortical LFP signatures of respiration and gamma are mediated by bulbar inputs in the form of a ROR, the bulk entrainment of limbic neuronal activity by breathing is mediated by an intracerebral RCD originating in the brainstem rhythm generators and being unaffected by olfactory de-afferentiation. We suggest that this joint modulation of limbic circuits by respiration is analogous to the predictive signaling employed in a wide range of neural circuits^64^, such as those underlying sensory-motor coordination^65^. Although the pathway of RCD remains unknown, we speculate that ascending long-range somatostatin-expressing interneuron projections from the preBötzinger complex to the thalamus, hypothalamus and basal forebrain^66^ or the locus coeruleus^67^ are probable pathways for this widespread modulation. The global and powerful nature of the RCD calls for future tracing and activity-dependent labeling studies to identify its anatomical substrate. A disinhibition-mediated mechanism of RCD would be consistent with the lack of prominent LFP sources in the absence of ROR, similar to the mechanism of disinhibitory pacing by the medial septum of the entorhinal-hippocampal system during theta oscillations^5^. We predict that the functional role of RCD in coordinating activity across the limbic system during offline states extends to other brain structures and brain states. The global outreach of RCD to higher order areas suggests that it might play an important role in the coordination of multisensory processing, in sync with orofacial motor output during both passive and active orofacial sampling, thus providing a centrifugal component synchronized with reafferent sensory inputs and respiratory efference copies to orofacial motor centers^68^.

We provide causal evidence that fear-related 4Hz oscillations^27,69^ are a state-specific expression of the limbic respiratory entrainment and originate from the reafferent respiratory entrainment of olfactory sensory neurons by passive airflow^35^. Importantly, although prefrontal 4Hz LFP oscillations originate in fear-associated enhanced breathing, the ROR is not necessary for the expression of innate or conditioned fear behavior, in agreement with a recent report^23^. This suggests the potential sufficiency of RCD for the behavioral expression. Interestingly, the optogenetic induction of such oscillations is sufficient to drive fear behavior in naïve animals^27^, raising the possibility that this effect is mediated by the bidirectional interaction of prefrontal networks with the RCD via top-down projections to PAG. This sets the stage for future investigations of the interaction between the RCD and ROR in limbic networks and in turn the top-down modulation of breathing and emotional responses.

We extend previous work on the hippocampal entrainment^21,22,38^ and provide new evidence for the mechanistic underpinnings of CA1 and DG modulation, by means of joint RCD and ROR inputs. We demonstrate robust modulation of hippocampal ripple occurrence by breathing, in agreement with a previous report^61^, and its effect on the response of prefrontal^50,70^ and accumbens neurons^71^ to SWR. Importantly, we provide the first evidence for the role of this global limbic circuit modulation by the respiratory rhythm in coordinating the interaction between the hippocampus and the downstream structures (i.e. mPFC and NAc), thought to underlie memory consolidation^72,73^.

Our results suggest that the respiratory rhythm orchestrates the hippocampo-cortical dialogue, by jointly biasing the neuronal firing and temporally coupling network dynamics across regions. We report for the first time the modulation of prefrontal UP and DOWN states as well as dentate spikes by breathing, which suggests a novel potential mechanism for the large-scale entrainment of thalamocortical excitability during sleep^74^. Along these lines, an OB-mediated pacing of slow oscillations in olfactory cortices has been demonstrated in the past^58^. Importantly, prefrontal SO entrainment appears to be dominated by the intracerebral RCD, while still receiving synchronous olfactory inputs via the ROR pathway. This highlights olfaction as a royal path to the sleeping brain that via synchronous ROR input reaches the limbic system in sync with RCD-coordinated UP-DOWN state dynamics during slow-wave sleep and could explain the efficacy of manipulations that bias learning^75^, consolidation^76^ or sleep depth^77^ using odor presentation during sleep. The intrinsic RCD-mediated comodulation of both SWR and SO by the respiratory phase brings into perspective the mechanistic explanation of studies that improve consolidation using stimulation conditioned on the ongoing phase of the cortical SO^15,78,79^. Understanding the causal role of respiratory entrainment in the formation, consolidation, and retrieval of memories will require fine time scale closed-loop optogenetic perturbation.

While a causal model of the role of respiration in temporally coordinating hippocampal ripples, dentate spikes, entorhinal cortex inputs, and prefrontal UP and DOWN states remains to be elucidated, their temporal progression and phase relationship with the ongoing respiratory cycle hints to a possible mechanism. ROR-driven gamma-associated waves and RCD lead to differential recruitment of the entorhinal cortex, consistent with the sinks in the dentate molecular layer and the entrainment of dentate spikes. Depending on the strength of the inputs, either feed-forward inhibition of the CA3^80^ or excitation during UP states^14^ can suppress or promote respectively SWRs. In parallel, the ripple-driven recruitment of prefrontal neurons likely triggers the resetting of the ongoing UP states by tilting the bias between excitation and inhibition^81^ and results in a feedback re-entrance to the entorhinal-hippocampal network^14^. Further analysis of the SO dynamics across the cortical mantle and their relationship with hippocampal ripples, as well as causal manipulation of the two nonlinear dynamics is required to validate and elucidate the physiological details of this model.

While we show here that the respiratory dynamics bias the prefrontal SO via ROR and RCD, slow oscillations can emerge in isolated cortical slubs^82^ or slices^83^. Furthermore, from a mechanistic perspective, the generative mechanism of the two oscillations is potentially comparable. Leading models of the generation of neocortical UP states from DOWN states^84^ or inspiratory bursts from expiratory silence in preBötzinger circuits^85^ suggest that both phenomena rely on regenerative avalanches due to recurrent connectivity, that are followed by activity-dependent disfacilitation. Given that neocortical slow oscillations can be locally generated^11,86^, are globally synchronized by the thalamic input^87^ and propagate across the neocortex^14,88^, ROR and RCD biasing of the cortical SO could be considered as an extension of a global system of mutually-coupled nonlinear oscillators. The persistent synchronous output of the respiratory oscillator and its marginal independence of the descending input might provide a widespread asymmetric bias to the slow oscillatory dynamics in the cortical circuits and SWR complexes in the hippocampus across offline states of different depth. It is likely, however, that via descending cortical projections, cortical SO provides feedback to the pontine respiratory rhythm-generating centers (e.g. via mPFC projections to PAG) and thus the interaction between respiratory dynamics and slow oscillations could be bidirectional.

This perpetual limbic rate comodulation by respiration also suggests a potential framework for memory-consolidation processes that do not rely on deep sleep and the associated synchronous K-complexes. This could explain the mechanism and distinctive role of awake replay in memory consolidation^89,90^. Given the substantial cross-species differences in respiratory frequency, as well as the effects of sleep depth and recent experience on cortical and hippocampal dynamics, it is conceivable that more synchronized and generalized UP states or awake vs. sleep ripples are differentially modulated by breathing. Further work is required to investigate the role of these parameters on the respiratory biasing of network dynamics and its role in memory consolidation.

Finally, in light of the wide modulation of limbic circuits by breathing during quiescence, we suggest that breathing effectively modulates the default mode network (DMN). To examine this hypothesis future work will be needed to carefully examine the fine temporal structure of neuronal assemblies and their modulation by the RCD and ROR copies of the breathing rhythm throughout cortical and subcortical structures, an endeavor that might uncover functional sub-networks of the DMN.

In summary, the data provided here suggest that respiration provides a perennial stream of rhythmic input to the brain. In addition to its role as the *condicio sine qua* non for life, we provide evidence that breathing rhythm acts as a global pacemaker for the brain, providing a reference signal that enables the integration of exteroceptive and interoceptive inputs with the internally generated dynamics of the limbic brain during offline states. In this emergent model of respiratory entrainment of limbic circuits, this common reference acts in tandem with the direct anatomical links between brain regions to pace the flow of information.

## Acknowledgments

We thank G. Schwesig, E. Blanco Hernandez and E. Resnik for valuable input, R. Ahmed for technical assistance, J. Lu for assistance in the experiments and all the members of the Sirota laboratory for helpful discussions and comments on the manuscript. This work was supported by grants from Munich Cluster for Systems Neurology (SyNergy, EXC 1010), Deutsche Forschungsgemeinschaft Priority Program 1665 and 1392 and Bundesministerium für Bildung und Forschung via grant no. 01GQ0440 (Bernstein Centre for Computational Neuroscience Munich) and European Union Horizon 2020 FETPOACT program via grant agreement no.723032 (BrainCom) (A.S.).

## Author Contributions

N.K and A.S. designed the experiments and data analysis, interpreted the data and wrote the manuscript, N.K. performed the experiments and analyzed the data.

## Methods

### Animals

Naive male C57BL6/J mice (3 months old, Jackson Laboratory) were individually housed for at least a week before all experiments, under a 12 light-dark cycle, ambient temperature 22 °C and provided with food and water ad libitum. Experiments were performed during the light phase. All procedures were performed in accordance with standard ethical guidelines (European Communities Directive 86/60-EEC) and stipulations of the German animal welfare law (Tierschutzgesetz ROB-55.2-2532.Vet_02-16-170). All efforts were made to minimize the number of animals used and the incurred discomfort.

### Surgery

Anesthesia was induced with 4% Isoflurane and surgical plane of anesthesia was maintained using 1% Isoflurane in *O*^2^. Body temperature was maintained at 37 °C with a custom heating pad. Analgesia was provided by means of subcutaneous administration of meloxicam (5 mg/kg) pre- and for 7 days post-operatively and local subcutaneous administration of a mixture of lidocaine (5 mg/kg) and bupivacaine (5 mg/kg). For free behavior recordings, electrode bundles, multi-wire electrode arrays or silicon probes were implanted chronically. Recordings targeted the medial prefrontal cortex (stereotaxic coordinates: 1.7-2 mm anterior to the bregma (AP), 0.3 mm lateral to the midline (ML) and 0.8 to 1.4 mm ventral to the cortical surface (DV)), dorsal hippocampus (AP: −2.3 mm, ML: 1.5 mm, DV: 0.8-1.5 mm), BLA (AP: −1.7 mm, ML: 3 mm, 4 mm DV) and NAc (AP: 1.2 mm, ML: 1 mm, DV: 4 mm)^91^. For head-fixed recordings, a craniotomy above the targeted structure and a midline bilateral craniotomy above the mPFC was performed to enable the recording from all cortical layers. Dura was left intact and craniotomies were sealed with Kwik-Cast (WPI, Germany) after surgery and after each recording session. For electromyographic (EMG) and electrocardiographic (ECG) recordings, two 125μm Teflon-coated silver electrodes (AG-5T, Science Products GmBH) were sutured into the right and left nuchal or dorsal intercostal muscles, using bio-absorbable sutures (Surgicryl Monofilament USP 5/0). Wires were connected to a multi-wire electrode array connector (Omnetics) attached to the skull. For the recording of the neural activity of the olfactory epithelium, which was used as a proxy for respiration^26^, a small hole was drilled above the anterior portion of the nasal bone (AP: +3 mm from the nasal fissure, ML: +0.5 mm from midline) until the olfactory epithelium was revealed. A 75μm Teflon-coated silver electrode (AG-3T, Science Products GmBH) was inserted inside the soft epithelial tissue. Approximately 500μm of insulation was removed from the tip of this wire and the other end was connected to the same Omnetics connector as the rest of the electrodes. Two miniature stainless steel screws (#000–120, Antrin Miniature Specialties, Inc.), pre-soldered to copper wire were implanted bilaterally above the cerebellum and served as the ground for electrophysiological recordings and as an anchoring point for the implants. All implants were secured using self-etching, light-curing dental adhesive (OptiBond All-In-One, Kerr), light-curing dental cement (Tetric Evoflow, Ivoclar Vivadent) and autopolymerizing prosthetic resin (Paladur, Heraeus Kulzer).

### Behavior

Electrophysiological recordings of the mice took place before and after each behavioral session in the home-cage. Home-cage consisted of clear acrylic filled with wood chip bedding and a metal grid ceiling which was removed for the purposes of the recordings. Food pellets were distributed in the home-cage and water was placed inside a plastic cup. Nesting material was available in the home-cage and utilized by the mice (typically building a nest in a corner). Exploratory behavior was recorded in a cheeseboard maze, consisting of a 60cm diameter acrylic cylinder with wooden laminated floor perforated with 10 mm diameter holes. For recordings of mice running freely on a wheel, a horizontal wheel (Flying Saucer) was permanently placed inside the homecage. The mice typically exhibited long running episodes on the wheel, with interspersed sleep episodes. Fear conditioning took place in context A, which consists of a square acrylic box (30 cm x 30 cm x 30 cm). Walls were externally decorated with black and white stripes. The box was dimly lit with white LEDs (25 lux) and white noise background sound was delivered through the walls using a surface transducer (WHD SoundWaver). The floor consisted of a custom designed metal grid (6 mm diameter stainless steel rods) connected to a precise current source (STG4004-1.6mA, Multi Channel Systems MCS GmbH). On day 1, mice were subjected to a habituation session in context A, during which the *CS*^+^ and *CS*^−^ (7.5 kHz, 80 dB or white-noise, 80 dB) were presented 4 times each. Each CS presentation consisted of 27 pips (50 ms duration, 2 ms rise and fall) with 1.1 s inter-pip interval. On the same day, in the fear conditioning session, *CS*^+^ was paired with the US. To serve as US, a mild electric foot-shock (1 s duration, 0.6 mA, 50 Hz AC, 5 CS-US pairings, 20–60 s randomized inter-trial intervals) was delivered to the mice through the metal grid. The onset of the foot-shock coincided with the offset of the conditioned stimulus. During the memory retrieval session, mice were presented with 4 *CS*^−^ and 4 *CS*^+^ presentations 24 hours after conditioning, in a distinct context B. For experiments involving pharmacological manipulation, a second retrieval session took place 12 days after fear conditioning. For experiments involving innate fear responses, mice were exposed for 10 min to a neutral context while a small filter paper, scented with the odorant 2-methyl-2-thiazoline (2MT) (M83406-25G; Sigma Aldrich), was placed in the environment. 2MT is a synthetic odorant, chemically related to the fox anogenital gland secretion 2,4,5-trimethyl-3-thiazoline (TMT), that induces robust innate fear responses, in contrast to TMT^92^. The sequence of the experimental protocol is schematized in Supplementary Fig. 3a.

### Behavioral analysis and state segmentation

A critical parameter for the behavior related analysis of neuronal activity is the proper determination of the behavioral state of the animal. For the purpose of behavioral state detection in freely-behaving mice, the movement of the animal was tracked using a 3-axis accelerometer (ADXL335, Analog Devices) incorporated in the headstage, which was used as the ground truth for the head-motion. Accelerometer data were sampled at 30 kHz and the sensitivity of the accelerometer is 340 mV/g (g is the standard acceleration due to gravity; ~ 9.8*m/s*^2^)). Since the accelerometer measures simultaneously the dynamic acceleration due to head movement and the static acceleration due to gravity, the first time derivative of the acceleration was calculated (jerk; units: g/s). This measure eliminates the effect of gravity and the dynamic acceleration dominates. The effect of gravity on the different axes is amplified during head rotations. The jerk of each axis was analyzed separately for the quantification of head-motion, however for the behavioral state detection the magnitude of the jerk was quantified as: 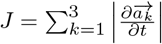, where 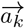 is the acceleration for each axis and smoothed in time using a narrow Gaussian window (2 s, 100 ms s.d.). Additionally, the activity of mice was tracked using an overhead camera (Logitech C920 HD Pro). The camera data were transferred to a computer dedicated to the behavior tracking and were acquired and processed in real-time using a custom-designed pipeline based on the Bonsai software^93^. Video data were synchronized with the electrophysiological data using network events. Video was preprocessed to extract the frame-to-frame difference and calculate a compound measure that we found provided an excellent proxy for the behavioral state. Complete immobility is easily distinguishable using this measure, due to the low amplitude and small variance. A threshold was set manually such that even small muscle twitches during sleep were captured, but breathing-related head-motion was below threshold. Using the density of head micro-motions and muscle twitches, we were able to classify behavioral segments as active awakening, quiescence or sleep (**Supplementary Fig. 1f**). For head-fixed recordings, we relied solely on high-resolution video of the mouse snout and body, from which we derived micromotion signal used in the same way as the jerk-based signal for freely-behaving mice.

### Head-fixed recordings

For high-density silicon probe recordings, we exploited the advantages of the head-fixed mouse preparation Mice were implanted with a lightweight laser-cut stainless steel headplate (Neurotar) above the cerebellum. After recovery from surgery, mice were habituated daily for 3-4 days to head-fixation prior to experimentation. A modified Mobile HomeCage (Neurotar) device was used, enabling mice to locomote, rest, and occasionally transition to sleep, within a customized free-floating carbon fiber enclosure (180 mm diameter and 40 mm wall height). Animal behavior was monitored using two modified 30fps, 1080p infrared cameras (ELP, Ailipu Technology Co), equipped with modified macro zoom lenses.

### In vivo electrophysiology

LFP and single-unit activity were recorded using either 12.5μm Teflon coated Tungsten wire (California Fine Wire) or custom-designed silicon probes (Neuronexus). High-density silicon probes (A1×64-Poly2-6mm-23s-160) were used for hippocampal CSD profiles, prefrontal depth profiles while multi-shank probes were used for prefrontal CSD analyses (A16×1-2mm-50–177). Individual electrodes or probe sites were electroplated to an impedance of 100-400 kΩ(at 1 kHZ) using a 75% polyethylene glycol - 25% gold^94^ or PEDOT solution^95^. NanoZ (White Matter) was used to pass constant electroplating current (0.1 – 0.5μA) and perform impedance spectroscopy for each electrode site. A reversed-polarity pulse of 1 s duration preceded the plating procedure to clean the electrode surface. After electroplating, electrodes impedance was tested in saline (at 1 kHz) and arrays were checked for shorts. Electrodes were connected to RHD2000 chip based amplifier boards (Intan Technologies) with 16–64 channels. Broadband (0.1 Hz-7.5 kHz) signals were acquired at 30 kHz. Signals were digitized at 16 bit and multiplexed at the amplifier boards and were transmitted to the OpenEphys recording controller using thin (1.8 mm diameter) 12-wire digital SPI (serial peripheral interface) cables (Intan Technologies). Typically 32-256 channels were recorded simultaneously. Data acquisition was synchronized across devices using custom-written network synchronization code. Breathing was measured using EOG recordings^26^. Following OD, the amplitude of the EOG signal was dramatically reduced. To quantify the effect of this manipulation on the neuronal entrainment by breathing, we recorded the respiratory rhythm using a fast response thermistor (GLS9-MCD, TE Connectivity) placed in close proximity to the naris of head-fixed mice.

### LFP analysis

Raw data were converted to binary format, low pass-filtered (0.5–400 Hz) to extract the local field potential component (LFP) and downsampled to 1 kHz. LFP signals were filtered for different frequency bands of interest using zero-phase-distortion sixth-order Butterworth filters. All data analysis was performed using custom-written software. Neuroscope data browser was used to aid with data visualization^96^.

### Spectral analysis

LFP power spectrum and LFP–LFP coherence estimations were performed using multitaper direct spectral estimates. For respiration frequency analyses, data were padded and a moving window of 3 s width and 2.4 s overlap was applied to the data. Signals were multiplied with 2 orthogonal taper functions (discrete prolate spheroidal sequences), Fourier transformed and averaged to obtain the spectral estimate^97^. For gamma frequency analyses, a window of 100 ms with 80 ms overlap, and 4 tapers were used. For some analyses and examples, to obtain a higher resolution in both time and frequency domain, data were transformed using complex Morlet wavelets (bandwidth parameter: 3, center frequency: 1.5). Convolution of the real and imaginary components of the transformed signal enables the extraction of the instantaneous amplitude and phase of the signal for each scale. For some example signal visualizations, we found it useful to utilize the real-part of the wavelet transformed signal, which preserves both phase and amplitude information (**Supplementary Fig. 6b**). For the power comodulation analysis^98^, the instantaneous multitaper estimate of the spectral power time series for each frequency bin in each structure was calculated and the Spearman correlation coefficient of every pair was calculated. To characterize the causal relationship between the respiratory signal and the prefrontal LFP, spectrally resolved Granger causality was calculated using the multivariate Granger causality toolbox^99^. Briefly, unfiltered LFP traces were detrended and normalized. The order of the vector autoregressive (VAR) model to be fitted was calculated using the Akaike information criterion.

### Phase modulation analysis

For phase analyses, the signal was filtered in the desired narrow frequency band and the complex-valued analytic signal was calculated using the Hilbert transform *ρ(**t**) = e^-iϕ(**t**)^*. The instantaneous amplitude at each timepoint was estimated based on the vector length, while the instantaneous phase of the signal was computed as the four-quadrant inverse tangent of the vector angle. A phase of 0° corresponds to the peak of the oscillation and a phase of 180° to the trough of the oscillation. The waveshape of the respiratory signal and its LFP counterparts are highly asymmetric, resulting in non-uniform phase distribution of this reference signal (**Supplementary Fig. 4c**). This deviation from uniformity is catastrophic for the phase modulation statistics since it biases the phase detection leading to false positive results. To account for this potential bias, the circular ranks of the phase distribution were computed and the phase distribution was transformed using the inverse of the empirical cumulative density function (ECDF) to return a signal with uniform prior distribution. After this correction, the phases can be assumed to be drawn from a uniform distribution enabling the unbiased application of circular statistics^8,27,41^. Point-processes with <200 events in the periods of interest were excluded from phase analyses, due to sample-size bias of these analyses^41^. For the quantification of phase modulation, the variance-stabilized log-transformed Rayleigh’s test Z 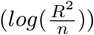, where Ris the resultant length and n the sample size, log is natural logarithm) was used^8,27,41^. This statistic quantifies the non-uniformity of a circular distribution against the von Mises distribution. Since ECDF transformation nonlinearly distorted phase axis circular mean of non-corrected phase samples were used for characterizing the preferred phase.

### Phase-amplitude cross-frequency coupling

For power-phase cross-frequency coupling, the modulation index (MI), as well as the mean resultant length (MRL), was calculated for each phase and amplitude pair^100^. Phase was evaluated for 1-20 Hz with a bandwidth of 1 Hz and step of 0.2 Hz using the Hilbert transform and correction for non-uniformity as described above. The amplitude was evaluated for 20-120 Hz with 5 Hz bandwidth and 3 Hz step. Shuffling statistics were used to evaluate the statistical significance of the MI and MRL by shuffling the phase and amplitude values.

### Current-source density analysis

Current-source density analysis was performed using the inverse CSD method^101^ with activity diameter 1 mm for slow and 0.5 mm for fast network events, 0.05 s.d., smoothed using varying cubic splines and extracellular conductivity *σ = 0.3S/m* based on calculations of isotropic and ohmic tissue impedance^102,103^. Importantly, all results were qualitatively confirmed by exploring the parameter space as well as using the classic second derivative CSD estimation method^104^. Occasional malfunctioning recording sites were interpolated from neighboring sites and all relevant sinks and sources were characterized and quantified from portions of data with no interpolated sites.

### Layer assignment

For the hippocampal high-density silicon probe recordings, channel layer assignment was performed based on established electrophysiological patterns of activity for different laminae^5^. The middle of the pyramidal layer was assigned to the channel with the highest amplitude of ripple oscillations (100-250 Hz band) and associated spiking activity. Neurons recorded dorsal of the channel with the highest SWR power were characterized as deep CA1 pyramidal neurons^40^. Conversely, neurons recorded ventral of this reference channel were characterized as superficial CA1 pyramidal neurons. Given that neuronal spikes can be identified in more than one channel of the polytrode, neurons were assigned to the channel with highest spike amplitude^105^. Well-described CSD profiles of hippocampal oscillatory patterns were used to assign somato-dendritic CA1 and DG layers to channels (**Supplementary Fig. 5d**). The middle of stratum radiatum was assigned to the channel with the deepest sharp wave current-source density sink associated with ripple oscillations^106,107^. Stratum oriens was defined as the channels above the pyramidal layer SWR CSD source and below the internal capsule, characterized by a positive component of the sharpwaves. For the identification of DG layers, we used the CSD and amplitude versus depth profile of dentate spikes (DS)^49^. DS are large amplitude events that occur naturally during offline states and reflect synchronized bursts of medial and lateral entorhinal cortex^49^. The outer molecular layer was defined as the Type-I dentate spike (DSI) sink, while the middle molecular layer was assigned as the channels exhibiting DSII sinks. The inner molecular layer was defined as the channel of the deepest secondary sink in the SWR triggered CSD, which is ventral of the DSII middle molecular layer sink. The source of DSII spikes, which corresponds to a typically more localized source preceding SWR events^107^, together with the polarity reversal of the DSII, which occurs above the granule cell layer^49^, enables the precise detection of this layer^108^. Stratum CA1 lacunosum-moleculare was defined as the difference between the theta-trough triggered CSD sink and the outer molecular DSII sink. This corresponds to approximately the dorsal third of the theta sink.

### Network event detection

Ripples were detected from a CA1 pyramidal layer channel using the instantaneous amplitude of the analytic signal calculated from the band-pass filtered (80-250 Hz). The instantaneous amplitude was referenced to the amplitude of a channel typically from the cortex overlying the hippocampus, was convolved with a Gaussian kernel (100 ms, 12 ms s.d.) and normalized. The mean and s.d. of the referenced amplitude were calculated for periods of quiescent immobility and slow-wave sleep. Ripples were detected as events exceeding 3 s.d with a minimum duration of 4 cycles and were aligned on the deepest trough of the bandpass filtered signal. Gamma bursts were detected using a similar procedure, but for the relevant frequency band and behavioral states. Dentate spikes (DS) were detected as large deviations (>3 s.d.) of the envelope of the 2-50 Hz band-pass filtered LFP signal from the DG hilar region, referenced to the CA1 pyramidal layer. Following detection, DS were clustered in two types using k-means clustering on the 2D space defined by the 2 principal components of the CA1/DG depth profile of each spike. UP and DOWN states were detected by binning the spike train for every single unit in 10 ms windows, normalized and convolved with a 0.5 s wide, 20 ms s.d. Gaussian kernel. The average binned spike histogram was calculated across all simultaneously recorded cells (including PNs and INs). DOWN states were detected as periods longer than 50 ms with no spikes across all the cells and the exact onset and offset of DOWN states were detected. UP states were detected as periods contained between two DOWN states, lasting between 100 ms and 2000 ms, with the average MUA activity during this period exceeding the 70th percentile of the MUA activity throughout the recording.

### Single-unit analysis and classification

Raw data were processed to detect spikes and extract singleunit activity. Briefly, the wide-band signals were band-pass filtered (0.6 kHz-6 kHz), spatially whitened across channels and thresholded and putative spikes were isolated. Clustering was performed using the ISO-SPLIT method implemented in MountainSort package^109^ and computed cluster metrics were used to pre-select units for later manual curation. Specifically, only clusters with low overlap with noise (<0.05), low peak noise (<30) and high isolation index (>0.9) were considered for manual curation, using custom-written software. At the manual curation step, only units with clean inter-spike interval (ISI) period, clean waveform, and sufficient amplitude were selected for further analysis. For the data collected with high-density polytrodes, after manual curation, a template of the spike waveform across 10 geometrically adjacent channels was calculated and the unit was re-assigned to the channel with largest waveform amplitude. To classify single-units into putative excitatory and inhibitory cell, a set of parameters based on the waveform shape, firing rate, and autocorrelogram were calculated. The two parameters that offered the best separation, in accordance to what has been reported in the past, were the trough-to-peak duration^28^ and the spike-asymmetry index (the difference between the pre- and post-depolarization positive peaks of the filtered trace)^110^, reflecting the duration of action potential repolarization which is shorter in interneurons^111,112^ (**Supplementary Fig. 4b**). Single-units with <200 spikes in the periods of interest were excluded from all analyses.

### Pharmacology

To causally prove the role of respiratory epithelium neurons in driving oscillations in the prefrontal cortex of mice, we induced a selective degeneration of the olfactory epithelium cells by systemic administration of methimazole^47^. Mice were injected intraperitoneally with methimazole (75 mg/kg). The effect on neuronal dynamics was characterized at 3, 7 or 10 days following the ablation of OSNs, with no appreciable differences between these timepoints.

### Statistical analysis

For statistical analyses, the normality assumption of the underlying distributions was assessed using the Kolmogorov-Smirnov test, Lilliefors test, and Shapiro-Wilk tests. Further, homoscedasticity was tested using the Levene or Brown-Forsythe tests. If the tests rejected their respective null hypothesis non-parametric statistics were used, alternatively, parametric tests were performed. When multiple statistical tests were performed, Bonferroni corrections were applied. Where necessary, resampling methods such as bootstrap and permutation tests were used to properly quantify significance. For box plots, the middle, bottom, and top lines correspond to the median, bottom, and top quartile, and whiskers to lower and upper extremes minus bottom quartile and top quartile, respectively.

### Anatomical analysis

After plating, electrodes and silicon probes were coated with DiI (ThermoFischer Scientific), a red fluorescent lipophilic dye^113^. Upon insertion in the brain, the dye is slowly incorporated in the cell membranes and diffuses laterally along the membrane, allowing the visualization of the electrode track and the histological verification of the electrode position. After the conclusion of the experiments, selected electrode sites were lesioned by passing anodal current through the electrode 0^114^. Typically, 10μ*A* current was passed for 5 s to produce lesions clearly visible under the microscope. One day was allowed before perfusion, to enable the formation of gliosis. Electrode tip locations were reconstructed with standard histological techniques. Mice were euthanized and transcardially perfused through the left ventricle with 4% EM grade paraformaldehyde (PFA) (Electron Microscopy Sciences) in 0.1 M PBS. Brains were sectioned using a vibratome 50 – 80μ*m* thick sections) and slices were stained with DAPI and mounted on gelatin-coated glass microscopy slides.

### Data availability

All relevant data that support the findings of this study will be made available from the corresponding author upon reasonable request.

### Code availability

Custom code used to acquire, process and analyze these data is available online (DataSuite, Nikolaos Karalis; https://github.com/nikolaskaralis/data_suite).

## Supplementary Figures

**Supplementary Figure 1.**
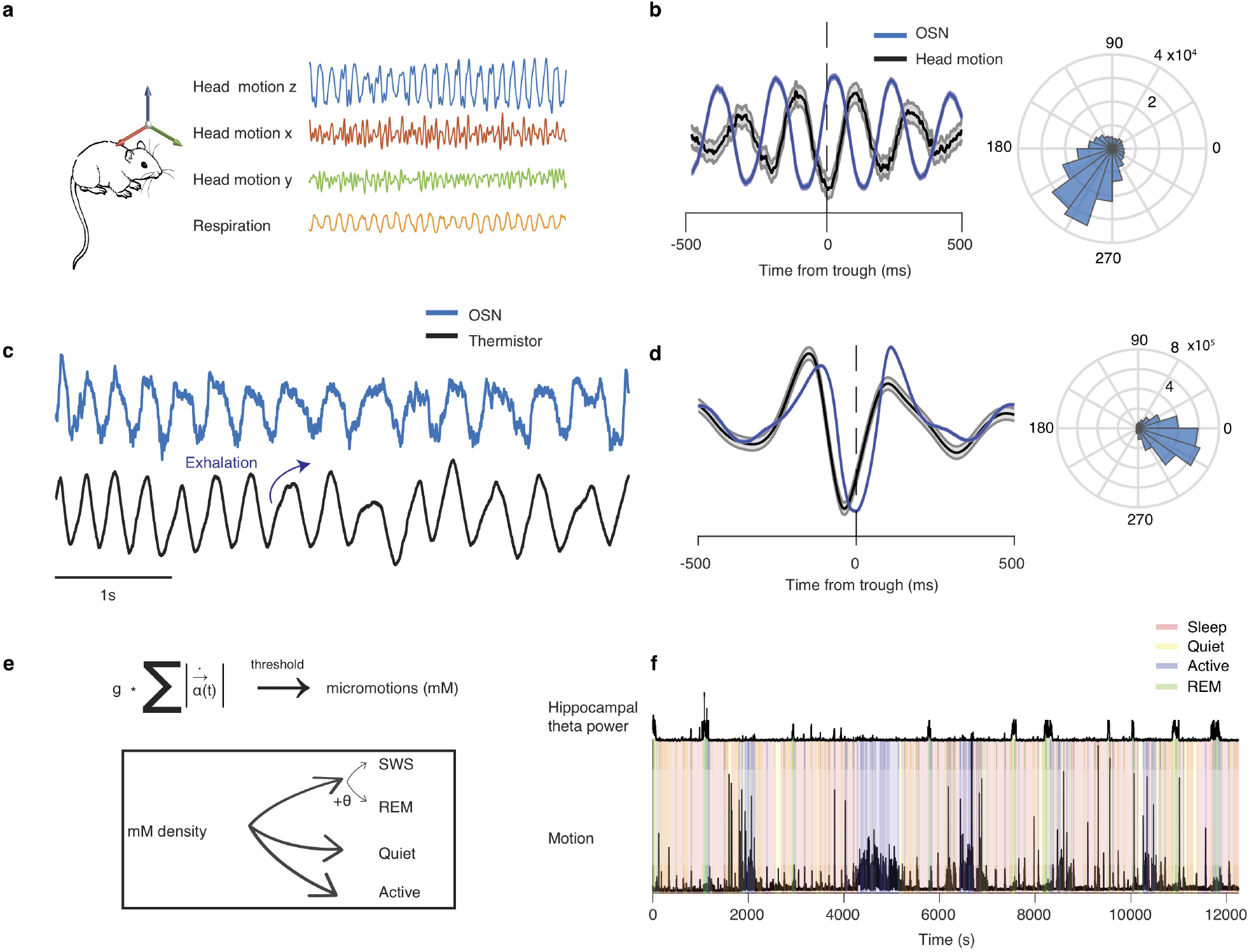
Validation of EOG recordings and behavioral state classification. (**a**) Schematic and example head-motion, respiration, and electrocardiogram (ECG) traces from freely-behaving mice. (**b**) Left, EOG trough-triggered EOG and head-motion signals during quiescence and sleep. Note the reliable phase relationship between the two signals. Right, distribution of phase shift between EOG and head-motion signals. (**c**) Example traces of EOG and simultaneous thermistor measurement. (**d**) EOG trough-triggered EOG and thermistor traces. Right, distribution of phase shift between the two signals. (**e**) Algorithm for the detection and classification of different behavioral states using behavioral variables extracted from the head-motion and based on the calculation of micro–motion (mM) density and hippocampal theta power. (**f**) Example classification of behavioral states from a recording in the home cage, using compound motion and hippocampal theta power. Shaded areas, mean ± s.e.m.

**Supplementary Figure 2.**
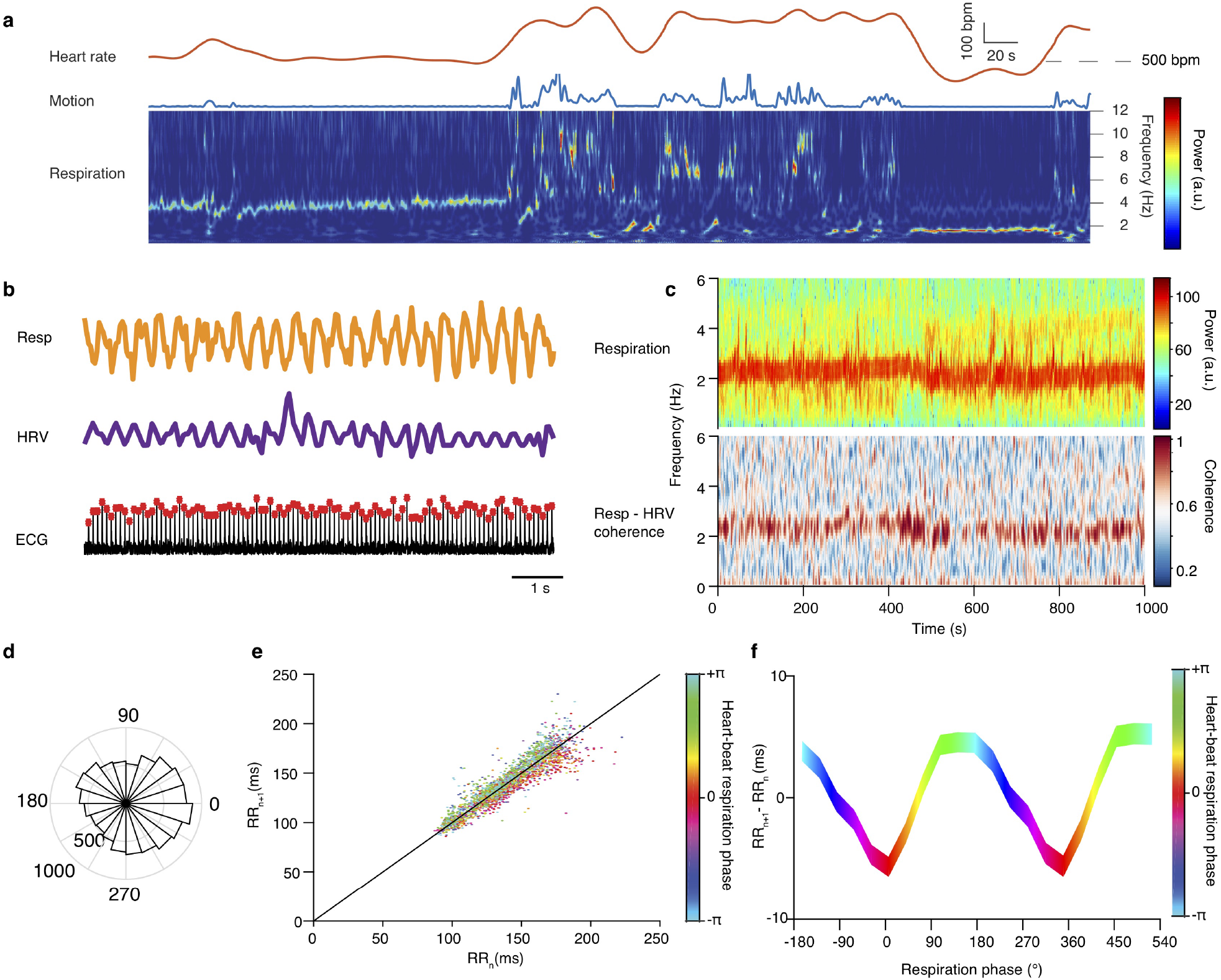
State-dependent cardiac - respiratory dynamics. (**a**) Example heart rate and compound motion traces (top) and spectral decomposition of respiration (bottom), from a quiet and sleeping mouse, demonstrating the intricate relationship between heart rate and respiratory frequency during different behavioral states. Dashed horizontal line marks the 500 bpm level. (**b**) Example respiration, heart-rate variability and raw electrocardiogram signals. (**c**) Example spectrogram of respiration (top) and coherogram between respiration and heart-rate variability (bottom). Note the almost perfect coherence between the two signals, highlighting the powerful effect of respiration in modulating the heart rhythm. (**d**) Distribution of the respiratory phase of each heart-beat reveals a phase preference. (**e**) Poincaré return map between consecutive R-R intervals. R-R intervals are color-coded based on the concurrent respiratory phase. The post-inspiratory events deviating from the diagonal are contributing to the observed relation between heart-rate variability and respiration. (**f**) Color-coded average time difference (variability) between consecutive R-R intervals as a function of the respiratory phase.

**Supplementary Figure 3.**
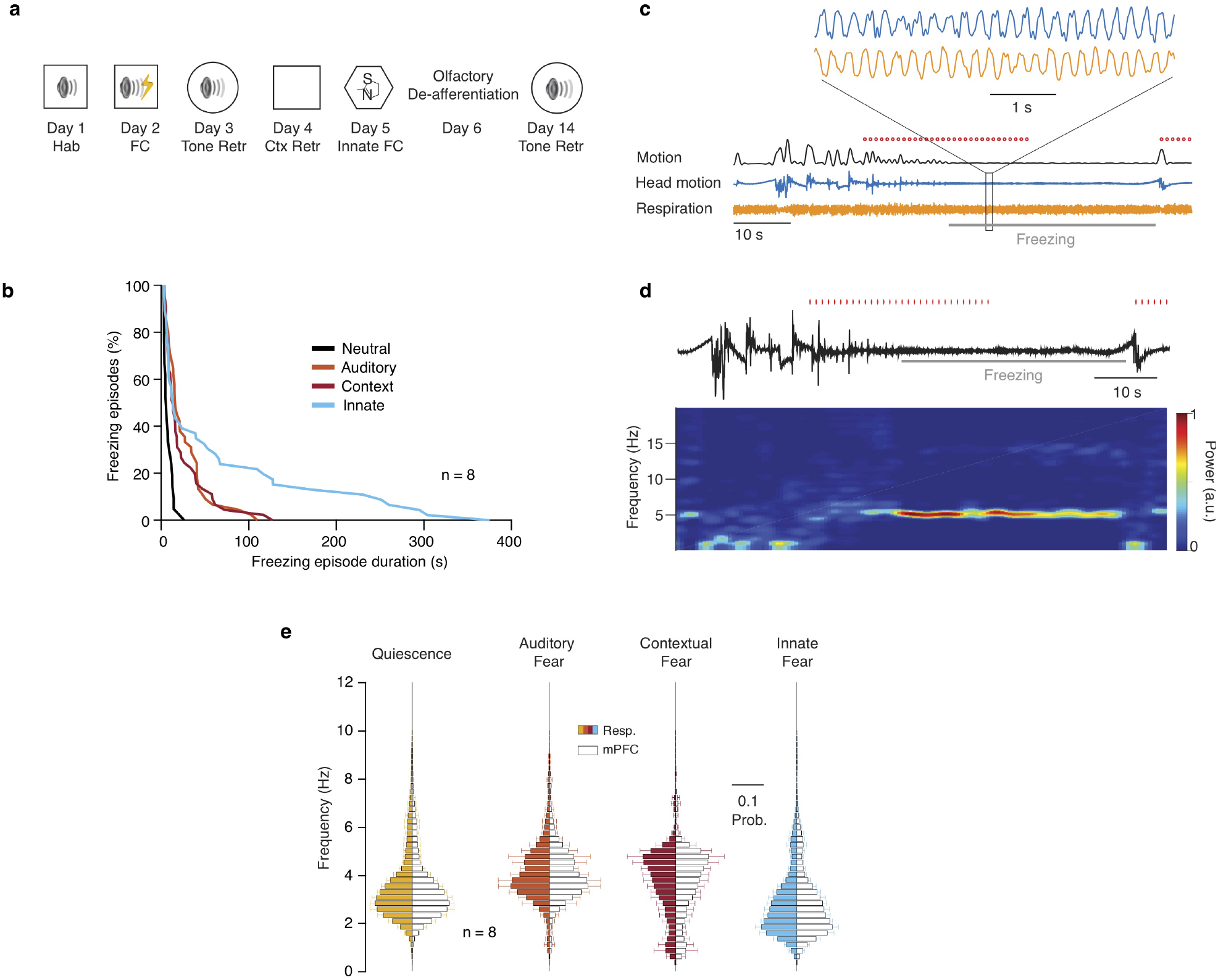
Respiratory dynamics during fear behavior. (**a**) Experimental protocol. (**b**) Cumulative distribution of freezing episode duration during neutral context exploration (black trace) and different types of fear behavior for all animals (n = 8 mice). (**c**) Example compound motion, head-motion and respiration traces during tone retrieval (day 3). Red ticks indicate CS presentations and the gray line indicates freezing episode duration. Inset, magnified traces during freezing reveals the presence of ~4 Hz respiration-related oscillatory patterns in EOG and head-motion. (**d**) Example head-motion trace (top) and spectral decomposition of this signal (bottom) during freezing behavior, revealing the presence of strong oscillatory components during freezing behavior. Red ticks indicate CS presentations and gray line indicates freezing episode duration. (**e**) Distribution of peak frequency bins of the spectrally decomposed respiration (left; darker colors) and mPFC LFP (right; lighter colors) during quiescence, auditory fear retrieval, contextual fear retrieval and innate fear (n = 8 mice). Hab., habituation; FC, fear conditioning; Ret., retrieval; Ctx Retr., Context retrieval.

**Supplementary Figure 4.**
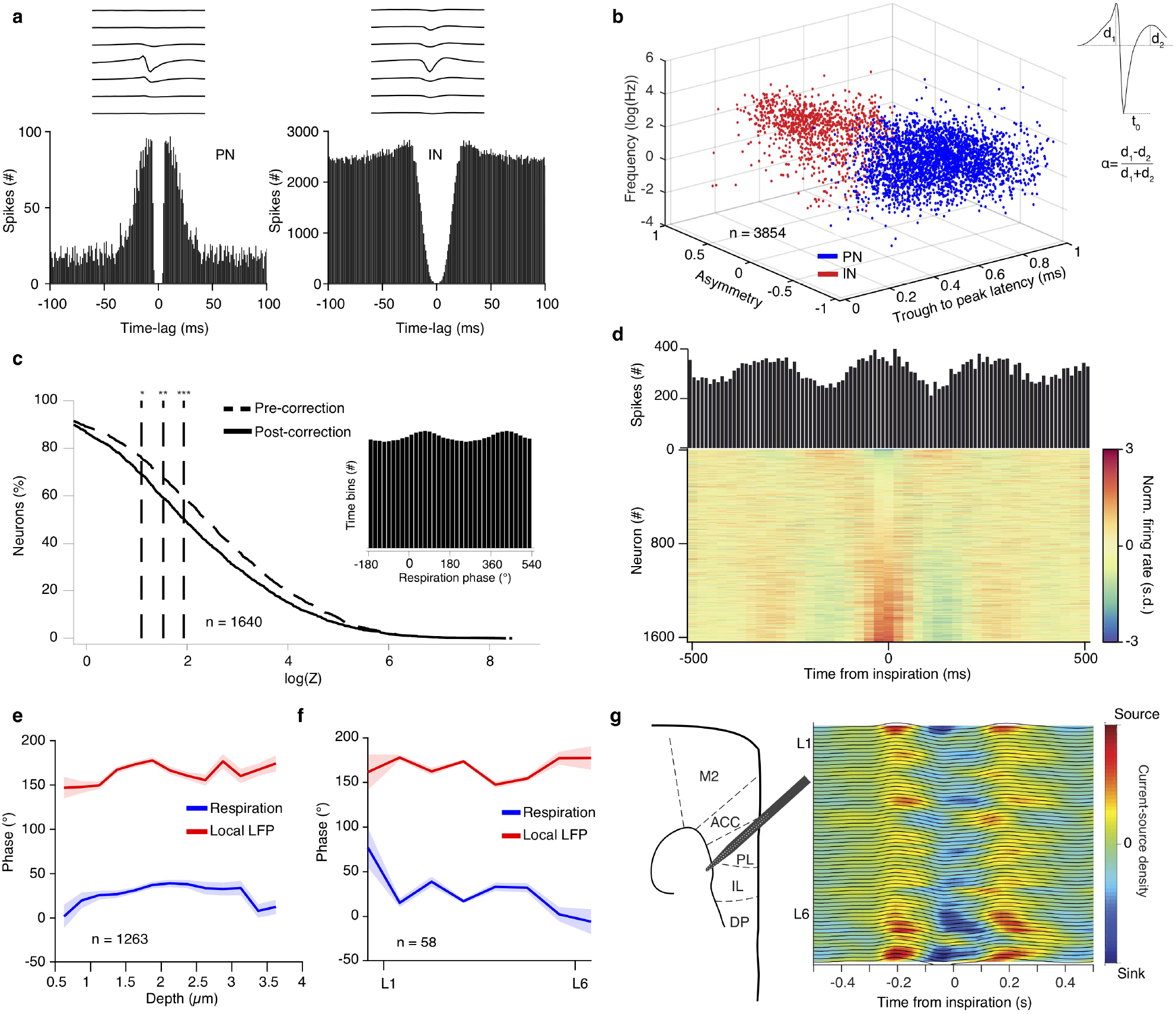
Unit classification and respiratory modulation. (**a**) Average spike spatio-temporal waveforms (top) and auto-correlation histograms (bottom) for an example prefrontal PN (left) and IN (right). (**b**) Neurons from all structures were classified as putative principal cells and interneurons using an unsupervised clustering method based on the firing frequency, the trough to peak latency (t_0_) and the waveform asymmetry index (**a**). (**c**) Cumulative distribution of the modulation strength (logZ) for all prefrontal neurons (n = 1640 cells), before and after correction for the non-uniformity of the respiratory phase. Uncorrected values are artifactually higher. Inset, example non-uniform prior distribution of the respiratory phase. (**d**) Top, example inspiration-triggered time histogram for an example prefrontal neuron. Bottom, color-coded normalized inspiration-triggered time histograms for all prefrontal neurons (n = 1640 cells), ordered by increasing normalized firing rate. (**e**) Depth-resolved average preferred phase for all prefrontal neurons (n = 1263 cells; n = 11 mice), assessed in relation to the phase of the respiration and the phase of the local LFP. (**f**) Translaminar average preferred phase for all prefrontal neurons (n = 58 cells; n = 6 mice), assessed in relation to the phase of the respiration and the phase of the local LFP. (**g**) Left, schematic of a single-shank angled high-density polytrode inserted in the dorsal mPFC. Right, Example high-resolution inspiration-triggered CSD of the mPFC respiration-related oscillation using a single-shank angled polytrode used to confirm the observations based on data acquired with the multi-shank silicon probe shown in Fig. 2j,k. Shaded areas, mean ± s.e.m. Stars indicate significance levels (* P<0.05; ** P<0.01; *** P<0.001). s.d., standard deviations. M2, motor area 2; ACC, anterior cingulate cortex; PL, prelimbic; IL, infralimbic; DP, dorsal peduncular; L1, layer 1; L6, layer 6.

**Supplementary Figure 5.**
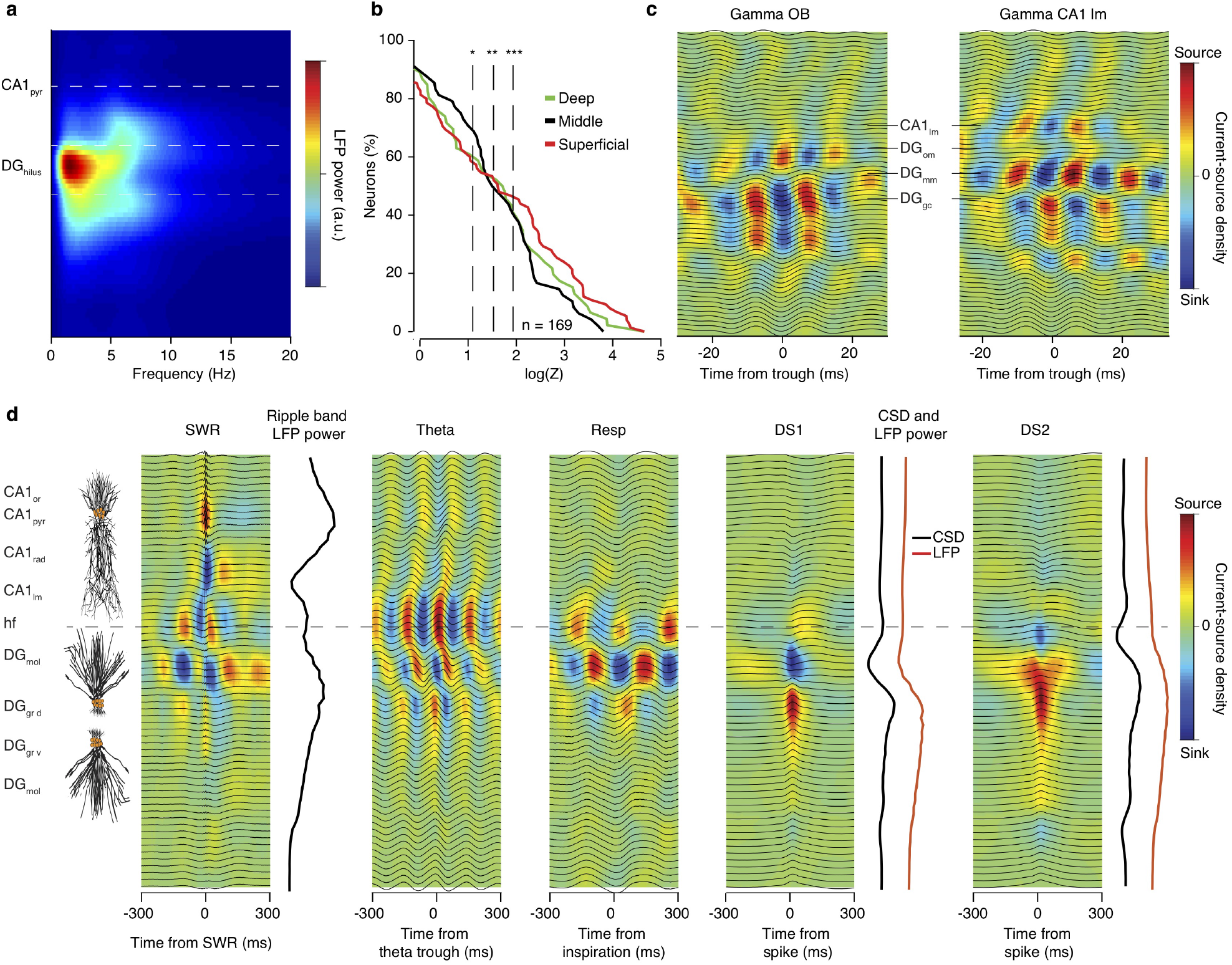
Translaminar profile of hippocampal dynamics. (**a**) Example translaminar profile of spectral power of the hippocampal LFP during quiescence and slow-wave sleep. Note strong peak in 2-4 Hz power reflecting DG LFP entrained by respiration (**b**) Cumulative distribution of the respiratory phase modulation strength for all CA1 putative pyramidal cells (n = 169 neurons), grouped based on their location within the pyramidal layer into deep (n = 46 cells), middle (n = 48 cells) and superficial (n = 75 cells) cells. (**c**) Example olfactory bulb gamma- and CA1lm gamma-triggered LFP traces and translaminar CSD in the dorsal hippocampus. (**d**) Left, schematic depiction of the neuronal alignment of CA1 pyramidal cells and DG granule cells, aligned to the CSD profiles as described in Methods section. Right, example CSD profiles of the dorsal hippocampus of the same animal, triggered on the peak of sharp-wave ripple (SWR) events, the troughs of theta oscillations, inspiratory events and the two types of dentate spikes (DS1 and DS2). The power of the ripple-band and the CSD magnitude and LFP power for the two types of dentate spikes is aligned to the CSD profiles. Horizontal dashed line indicates the hippocampal fissure. Stars indicate significance levels (* P<0.05; ** P<0.01; *** P<0.001). *CA1_or_*, oriens; *CA1_pyr_*, str. pyramidale; *CA1_rad_*, str. radiatum; *CA1_lm_*, str. lacunosum-moleculare; hf, hippocampal fissure; *DG_mol_*, dentate gyrus – molecular layer; *DG_om_*, outer molecular layer; *DG_mm_*, middle molecular layer; *DG_grd/v_*, granule cell layer (dorsal/ventral); *DG_hil_*, hilus.

**Supplementary Figure 6.**
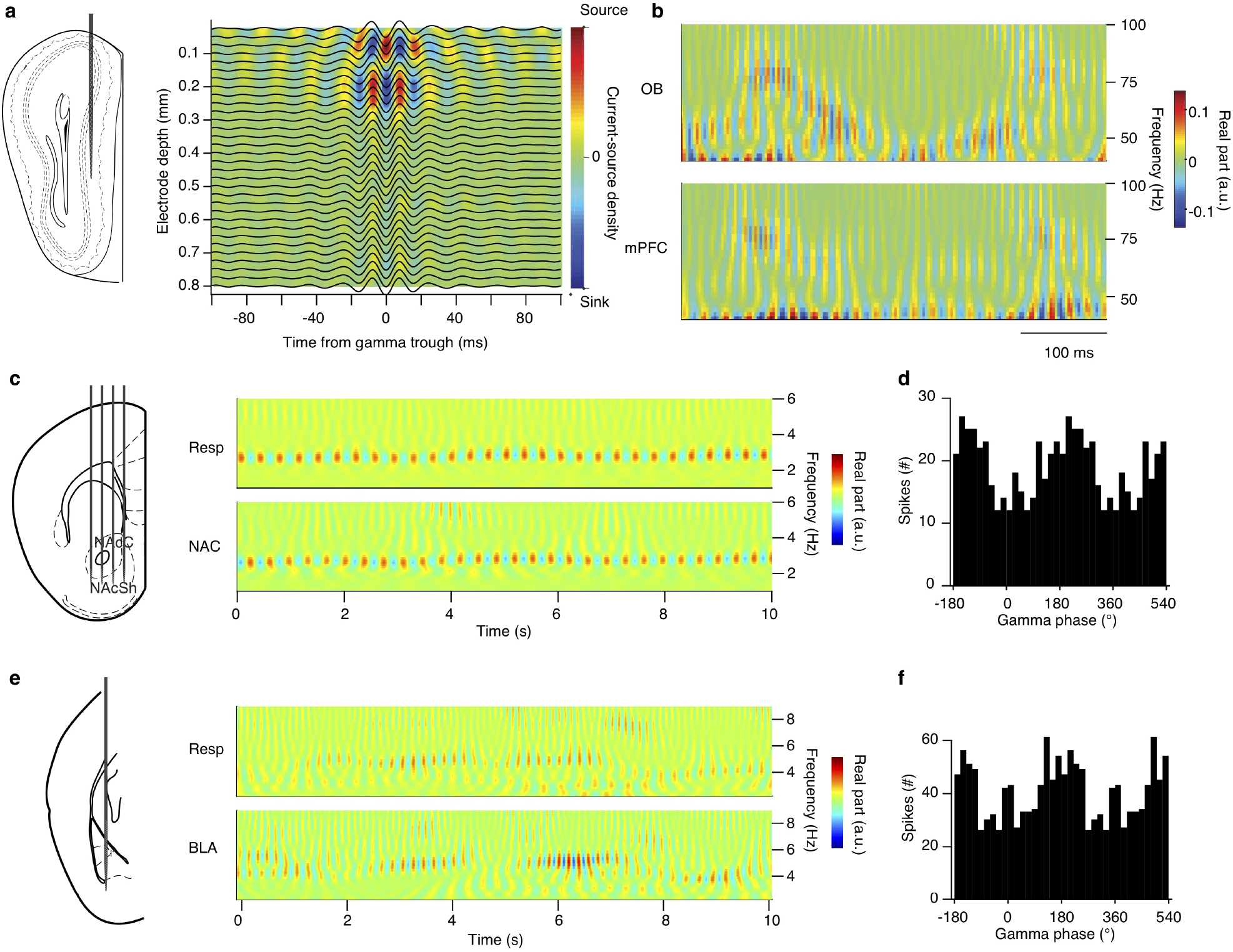
Gamma dynamics. (**a**) Left, schematic of a high-density polytrode recording across layers of the olfactory bulb. Right, example high-resolution inspiration-triggered current-source density of the olfactory bulb. (**b**) Example spectral representation of the OB and mPFC gamma-band LFP using the real-part of the wavelet transform highlights the transient phase relation between bursts of gamma oscillations in the two structures. (**c**) Left, schematic of a recording from the NAc using a 4 shank silicon polytrode. Right, example spectral representation of the respiration and NAc LFP using the real-part of the wavelet transform highlights the reliable phase relationship between the two signals. (**d**) Gamma phase distribution of the spikes of an example NAc neuron. (**e**) Left, schematic of a high-density polytrode recording from the BLA. Right, example spectral representation of the respiration and BLA LFP using the real-part of the wavelet transform highlights the reliable phase relationship between the two signals. (**f**) Gamma phase distribution of the spikes of an example BLA neuron. a.u., arbitrary units.

**Supplementary Figure 7.**
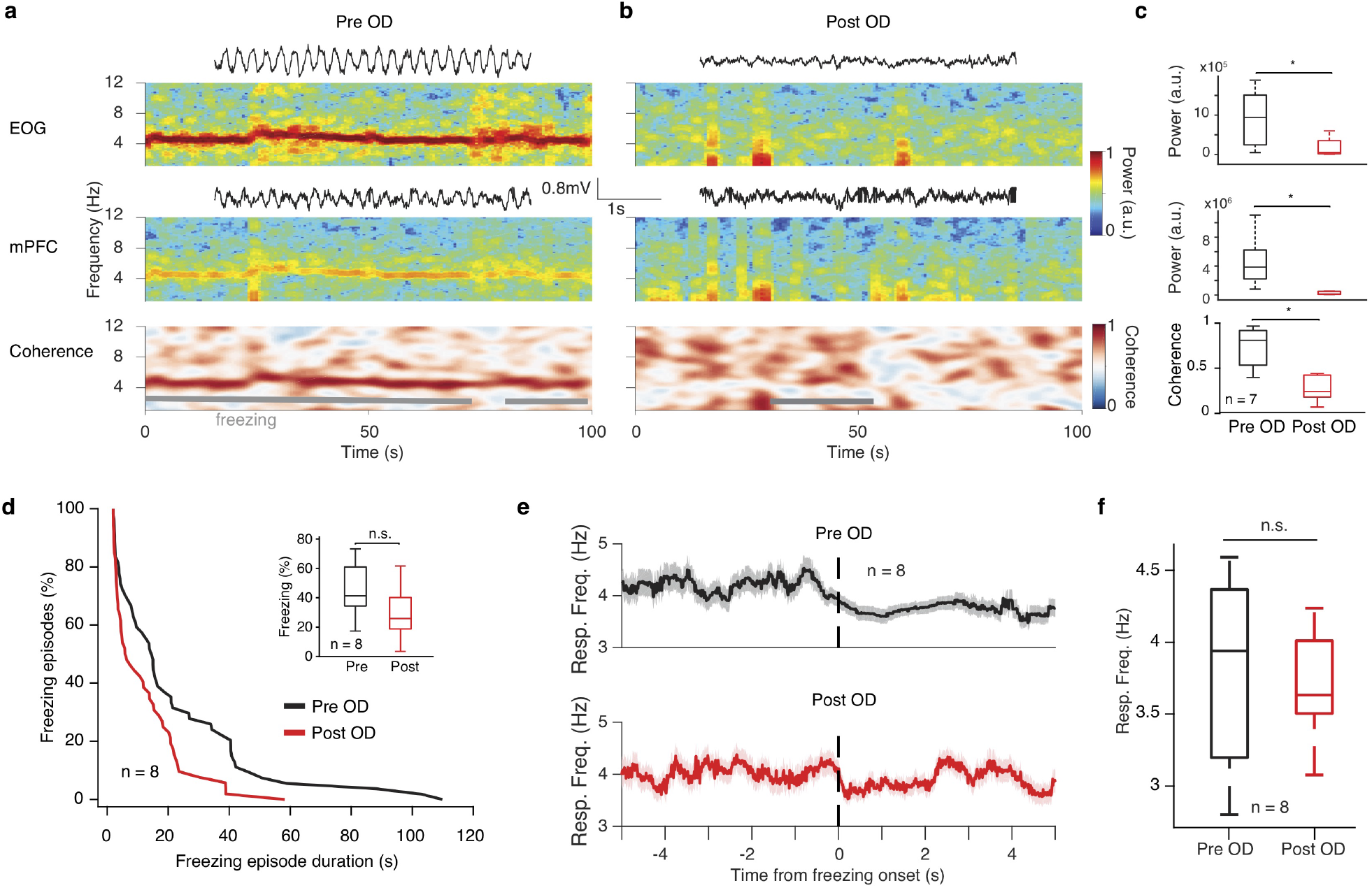
Effect of olfactory de-afferentiation on fear behavior. (**a**, **b**) Example spectral decomposition of the respiration (top) and mPFC LFP (middle) during freezing behavior and coherogram between the two signals (bottom), in baseline conditions (**a**) and after OD (**b**). (**c**) Average EOG (top) and mPFC LFP(middle) power and coherence between them (bottom) for the 2-5 Hz band before and after OD (n = 7 mice, Wilcoxon signed-rank tests in all plots, pre-versus post-OD). (**d**) Cumulative distribution of freezing episode duration for tone retrieval before and after OD (n = 8 mice). Inset, average freezing before and after OD (n = 8 mice, Wilcoxon signed-rank test, pre-versus post-OD, P = 0.109). (**e**) Freezing onset triggered respiratory frequency before and after OD (n = 8 mice). (**f**) Average respiratory frequency during freezing, before and after OD (n = 8 mice, Wilcoxon signed-rank test, pre-versus post-OD, P = 0.742). For box plots, the middle, bottom, and top lines correspond to the median, bottom, and top quartile, and whiskers to lower and upper extremes minus bottom quartile and top quartile, respectively. Stars indicate significance levels (* P<0.05; ** P<0.01; *** P<0.001). n.s., not significant; OD, olfactory de-afferentiation.

**Supplementary Figure 8.**
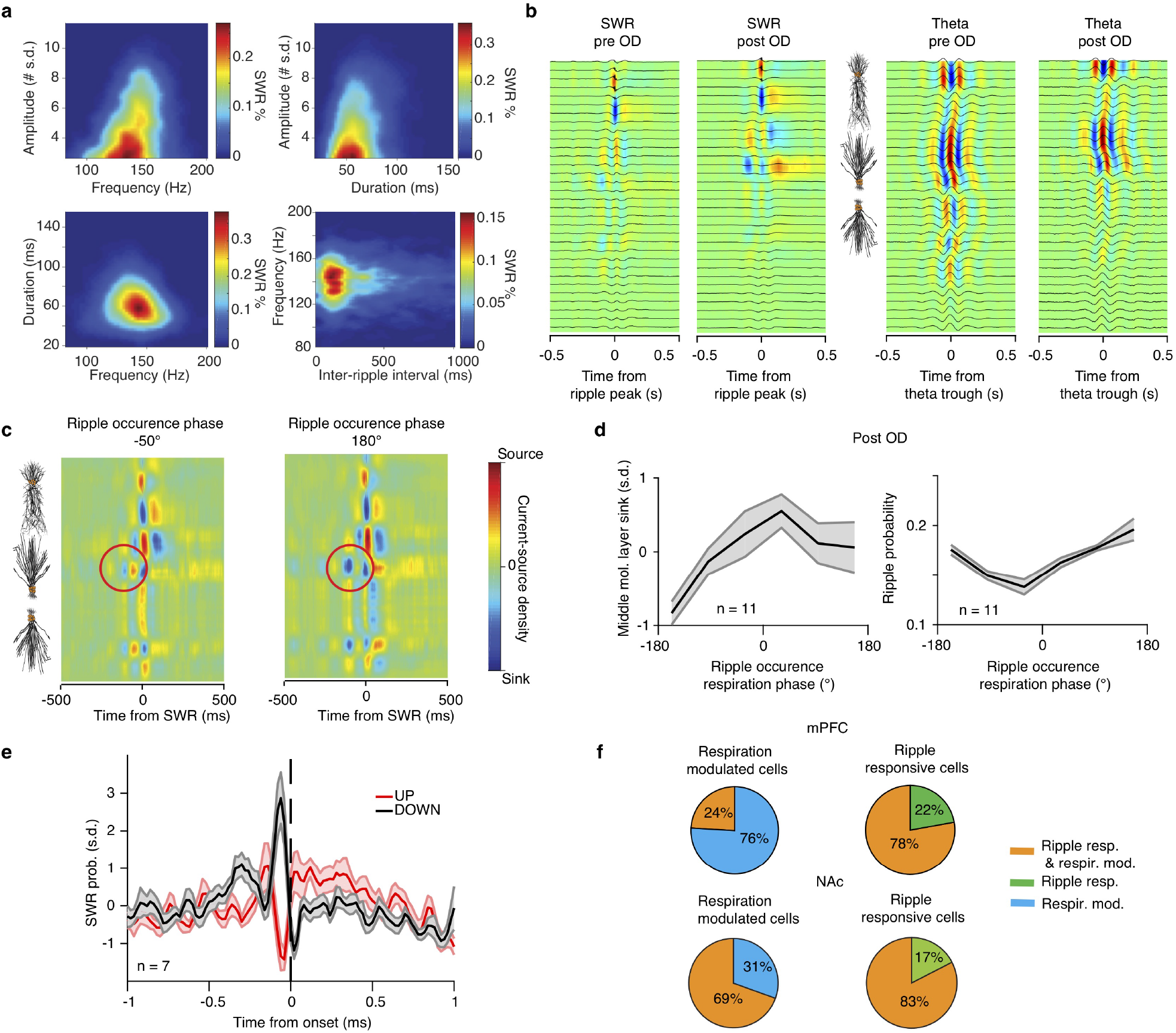
Effect of OD on characteristics of hippocampal respiration-related patterns. (**a**) Example color-coded pairwise joint probability distributions of the properties of ripple events, including the amplitude, frequency, duration and inter-ripple interval. (**b**) Example current-source density profiles of the dorsal hippocampus of the same animal, triggered on the peak of sharp-wave ripple (SWR) events and the troughs of theta oscillations (detected in CA1 pyramidal layer) before and after OD. (**c**) Example ripple-triggered translaminar current-source density profiles of the dorsal hippocampus grouped by the respiratory phase of the ripple occurrence: −50°(left) and 180°(right). Note strong sink at DG middle mol. Layer (red circle) present in the latter but not former. (**d**) Average respiration phase-resolved ripple-triggered middle molecular layer dentate sink magnitude (left) and corresponding phase ripple normalized incidence rate (right). (**e**) Probability of SWR occurrence as a function of time from UP or DOWN state onset after OD (n = 7 mice). Note that the observed pattern is identical to pre OD shown in Fig. 7i. (**f**) Pie charts indicating the percentage of mPFC neurons being either phase modulated by respiration (resp. mod), responding significantly to ripples (ripple resp.) or being both significantly modulated by respiratory rhythm and significantly responsive to ripples for all mPFC and NAc respiration-modulated cells and ripple-responsive cells. Shaded areas, mean ± s.e.m. Stars indicate significance levels (* P<0.05; ** P<0.01; *** P<0.001). OD, olfactory de-afferentiation.

## References

1. Papez, J. W. A proposed mechanism of emotions. Archives of Neurology & Psychiatry 38, 725–743 (1937).

2. Maclean, P. D. Psychosomatic disease and the visceral brain; recent developments bearing on the Papez theory of emotion. Psychosomatic medicine 11, 338–353 (1949).

3. Scoville, W. B. & Milner, B. Loss of recent memory after bilateral hippocampal lesions. Journal of Neurology, Neurosurgery & Psychiatry 20, 11–21 (1957).

4. Buzsaki, G. & Draguhn, A. Neuronal oscillations in cortical networks. Science 304, 1926–1929 (2004).

5. Buzsaki, G. Theta oscillations in the hippocampus. Neuron 33, 325–340 (2002).

6. Mizuseki, K., Sirota, A., Pastalkova, E. & Buzsaki, G. Theta oscillations provide temporal windows for local circuit computation in the entorhinal-hippocampal loop. Neuron 64, 267–280 (2009).

7. Fernández-Ruiz, A. et al. Entorhinal-CA3 dualinput control of spike timing in the hippocampus by theta-gamma Coupling. Neuron 93, 1213–1226.e5 (2017).

8. Siapas, A. G., Lubenov, E. V. & Wilson, M. A. Prefrontal phase locking to hippocampal theta oscillations. Neuron 46, 141–151 (2005).

9. Benchenane, K. et al. Coherent Theta Oscillations and Reorganization of Spike Timing in the Hippocampal-Prefrontal Network upon Learning. Neuron 66, 921–936 (2010).

10. Steriade, M. M., McCormick, D. A & Sejnowski, T. J. Thalamocortical oscillations in the sleeping and aroused brain. Science 262, 679–685 (1993).

11. Sirota, A. & Buzsaki, G. Interaction between neocortical and hippocampal networks via slow oscillations. Thalamus and Related Systems 3, 245–259 (2005).

12. Wilson, M. & McNaughton, B. Dynamics of the hippocampal ensemble code for space. Science 261, 1055–1058 (1993).

13. Sirota, A., Csicsvari, J., Buhl, D. L. & Buzsaki, G. Communication between neocortex and hippocampus during sleep in rodents. Proceedings of the National Academy of Sciences 100, 2065–2069 (2003).

14. Isomura, Y. et al. Integration and Segregation of Activity in Entorhinal-Hippocampal Subregions by Neocortical Slow Oscillations. Neuron 52, 871–882 (2006).

15. Maingret, N., Girardeau, G., Todorova, R., Goutierre, M. & Zugaro, M. Hippocampo-cortical coupling mediates memory consolidation during sleep. Nature Neuroscience 19, 959–964 (2016).

16. Girardeau, G. et al. Selective suppression of hippocampal ripples impairs spatial memory. Nature Neuroscience 12, 1222–1223 (2009).

17. Rothschild, G., Eban, E. & Frank, L. M. A cortical – hippocampal – cortical loop of information processing during memory consolidation. Nature Neuroscience 20, 1–12 (2016).

18. Maviel, T., Durkin, T. P., Menzaghi, F. & Bontempi, B. Sites of neocortical reorganization critical for remote spatial memory. Science 305, 96–99 (2004).

19. Kitamura, T. et al. Engrams and circuits crucial for systems consolidation of a memory. Science 356, 73–78 (2017).

20. Ito, J. et al. Whisker barrel cortex delta oscillations and gamma power in the awake mouse are linked to respiration. Nature Communications 5, 3572 (2014).

21. Yanovsky, Y., Ciatipis, M., Draguhn, a., Tort, a. B. L. & Branka, k.J. Slow Oscillations in the Mouse Hippocampus Entrained by Nasal Respiration. Journal of Neuroscience 34, 5949–5964 (2014).

22. Lockmann, A. L.V., Laplagne, D. A., Leão, R. N. & Tort, A. B. L. A respiration-coupled rhythm in the rat hippocampus independent of theta and slow oscillations. Journal of Neuroscience 36, 5338–5352 (2016).

23. Moberly, A. H. et al. Olfactory inputs modulate respiration-related rhythmic activity in the prefrontal cortex and freezing behavior. Nature Communications (2018).

24. Zelano, C. et al. qNasal respiration entrains human limbic oscillations and modulates cognitive function. Journal of Neuroscience 36, 12448–12467 (2016).

25. Herrero, J. L., Khuvis, S., Yeagle, E., Cerf, M. & Mehta, A.D. Breathing above the brainstem: Volitional control and attentional modulation in humans. Journal of Neurophysiol-ogy 119, 145–159 (2018).

26. Ottoson, D. Analysis of the electrical activity of the olfactory epithelium. Acta physiologica Scandinavica. Supplementum 35, 1–83 (1955).

27. Karalis, N. et al. 4-Hz oscillations synchronize prefrontal-amygdala circuits during fear behavior. Nature Neuroscience19, 605–612 (2016).

28. Bartho, P. Characterization of neocortical principal cells and interneurons by network interactions and extracellular features. Journal of Neurophysiology 92, 600–608 (2004).

29. Biskamp, J., Bartos, M. & Sauer, J. F. Organization of pre-frontal network activity by respiration-related oscillations. Scientific Reports 7, 45508 (2017).

30. Hoover, W. B. & Vertes, R. P. Anatomical analysis of afferent projections to the medial prefrontal cortex in the rat. Brain Structure and Function 212, 149–179 (2007).

31. Herry, C. & Johansen, J. P. Encoding of fear learning and memory in distributed neuronal circuits. Nature Neuroscience 17, 1644–1654 (2014).

32. Adrian, E. D. Olfactory reactions in the brain of the hedgehog. The Journal of Physiology 100, 459–473 (1942).

33. Macrides, F & Chorover, S. L. Olfactory bulb units: activity correlated with inhalation cycles and odor quality. Science 175, 84–87 (1972).

34. Fukunaga, I., Herb, J. T., Kollo, M., Boyden, E. S. & Schaefer, A. T. Independent control of gamma and theta activity by distinct interneuron networks in the olfactory bulb. Nature Neuroscience 17, 1208–1216 (2014).

35. Grosmaitre, X., Santarelli, L. C., Tan, J., Luo, M. & Ma, M. Dual functions of mammalian olfactory sensory neurons as odor detectors and mechanical sensors. Nature Neuroscience 10, 348–354 (2007).

36. Buzsaki, G. Neural syntax: cell assemblies, synapsembles, and readers. Neuron 68, 362–385 (2010).

37. Vanderwolf, C. H. Hippocampal activity, olfaction, and sniffing: An olfactory input to the dentate gyrus. Brain Research 593, 197–208 (1992).

38. Nguyen Chi, V. et al. Hippocampal respiration-driven rhythm distinct from theta Oscillations in awake mice. Journal of Neuroscience 36, 162–177 (2016).

39. Valero, M. et al. in Nature Neuroscience 9, 1281–1290 (Nature Publishing Group, 2015).

40. Mizuseki, K., Diba, K., Pastalkova, E. & Buzsaki, G. Hippocampal CA1 pyramidal cells form functionally distinct sublayers. Nature Neuroscience 14, 1174–1183 (2011).

41. Sirota, A. et al. Entrainment of neocortical neurons and gamma oscillations by the hippocampal theta rhythm. Neuron 60, 683–697 (2008).

42. Stujenske, J. M., Likhtik, E., Topiwala, M. A. & Gordon, J. A. Fear and safety engage competing patterns of theta-gamma coupling in the basolateral amygdala. Neuron 83, 919–933 (2014).

43. Adrian, E. D. The electrical activity of the mammalian olfactory bulb. Electroencephalography and Clinical Neurophysiology 2, 377–388 (1950).

44. Freeman, J. A. & Nicholson, C. Experimental optimization of current source-density technique for anuran cerebellum. Journal of neurophysiology 38, 369–382 (1975).

45. Lepousez, G. & Lledo, P.-M. Odor discrimination requires proper olfactory fast oscillations in awake mice. Neuron 80, 1010–1024 (2013).

46. Tort, A. B., Brankačk, J. & Draguhn, A. Respiration-entrained brain rhythms are global but often overlooked. Trends in Neurosciences 41, 186–197 (2018).

47. Bergman, U, Ostergren, A, Gustafson, A.-L. & Brittebo, B. Differential effects of olfactory toxicants on olfactory regeneration. Archives of toxicology 76, 104–112 (2002).

48. Buzsaki, G. & Schomburg, E. W. What does gamma coherence tell us about inter-regional neural communication? Nature Neuroscience 18, 484–489 (2015).

49. Bragin, A, Jandó, G, Nádasdy, Z, van Landeghem, M & Buzsaki, G. Dentate EEG spikes and associated interneuronal population bursts in the hippocampal hilar region of the rat. Journal of neurophysiology 73, 1691–705 (1995).

50. Siapas, A. G. & Wilson, M. A. Coordinated interactions between hippocampal ripples and cortical spindles during slow-wave sleep. Neuron 21, 1123–1128 (1998).

51. Peyrache, A., Battaglia, F. P. & Destexhe, A. Inhibition recruitment in prefrontal cortex during sleep spindles and gating of hippocampal inputs. Proceedings of the National Academy of Sciences 108, 17207–17212 (2011).

52. Khodagholy, D., Gelinas, J. N. & Buzsaki, G. Learning-enhanced coupling between ripple oscillations in association cortices and hippocampus. Science 358, 369–372 (2017).

53. Hobson, J. A. Respiration and EEG synchronization in the frog. Nature 213, 988–989 (1967).

54. Busek, P & Kemlink, D. The influence of the respiratory cycle on the EEG. Physiological research 54, 327–333 (2005).

55. Yuan, H., Zotev, V., Phillips, R. & Bodurka, J. Correlated slow fluctuations in respiration, EEG, and BOLD fMRI. NeuroImage 79, 81–93 (2013).

56. Phillips, M. E., Sachdev, R. N. S., Willhite, D. C. & Shepherd, G. M. Respiration drives network activity and modulates synaptic and circuit processing of lateral inhibition in the olfactory bulb. Journal of Neuroscience 32, 85–98 (2012).

57. Kay, L. M. & Freeman, W. J. Bidirectional processing in the olfactory-limbic axis during olfactory behavior. Behavioral Neuroscience 112, 541–553 (1998).

58. Fontanini, A., Spano, P. & Bower, J. M. Ketamine–Xylazine-Induced Slow (?1.5 Hz) Oscillations in the Rat Piriform (Olfactory) Cortex Are Functionally Correlated with Respiration. Journal of Neuroscience 23, 7993–8001 (2003).

59. Zhong, W. et al. Selective entrainment of gamma subbands by different slow network oscillations. Proceedings of the National Academy of Sciences 114, 4519–4524 (2017).

60. Poe, G. R., Kristensen, M. P., Rector, D. M. & Harper, R. M. Hippocampal activity during transient respiratory events in the freely behaving cat. Neuroscience 72, 39–48 (1996).

61. Liu, Y., McAfee, S. S. & Heck, D. H. Hippocampal sharp-wave ripples in awake mice are entrained by respiration. Scientific reports 7, 8950 (2017).

62. Courtin, J., Karalis, N., Gonzalez-Campo, C., Wurtz, H. & Herry, C. Persistence of amygdala gamma oscillations during extinction learning predicts spontaneous fear recovery. Neurobiology of Learning and Memory 113, 82–89 (2014).

63. Van der Meer, M. A. Low and high gamma oscillations in rat ventral striatum have distinct relationships to behavior, reward, and spiking activity on a learned spatial decision task. Frontiers in Integrative Neuroscience 3, 9 (2009).

64. Crapse, T. B. & Sommer, M. A. Corollary discharge across the animal kingdom. Nature Reviews Neuroscience 9, 587–600 (2008).

65. Straka, H., Simmers, J. & Chagnaud, B. P. A new perspective on predictive motor signaling. Current Biology 28, R193–R194 (2018).

66. Yang, C. F. & Feldman, J. L. Efferent projections of excitatory and inhibitory preBötzinger complex neurons. Journal of Comparative Neurology 526, 1389–1402 (2018).

67. Yackle, K. et al. Breathing control center neurons that promote arousal in mice. Science 355, 1411–1415 (2017).

68. Moore, J. D. et al. Hierarchy of orofacial rhythms revealed through whisking and breathing. Nature 497, 205–210 (2013).

69. Dejean, C. et al. Prefrontal neuronal assemblies temporally control fear behaviour. Nature 535, 420–424 (2016).

70. Jadhav, S. P. P., Rothschild, G., Roumis, D. K. K. & Frank, L. M. M. Coordinated excitation and inhibition of prefrontal ensembles during awake hippocampal sharp-wave ripple events. Neuron 90, 113–127 (2016).

71. Pennartz, C. M. A. The ventral striatum in off-line processing: ensemble reactivation during sleep and modulation by hippocampal ripples. Journal of Neuroscience 24, 6446–6456 (2004).

72. Buzsaki, G. Two-stage model of memory trace formation: a role for “noisy” brain states. Neuroscience 31, 551–570 (1989).

73. Squire, L. R. & Alvarez, P. Retrograde amnesia and memory consolidation: a neurobiological perspective. Current opinion in neurobiology 5, 169–177 (1995).

74. Headley, D. B., Kanta, V. & Paré, D. Intra- and interregional cortical interactions related to sharp-wave ripples and dentate spikes. Journal of Neurophysiology 117, 556–565 (2017).

75. Arzi, A. et al. Humans can learn new information during sleep. Nature Neuroscience (2012).

76. Rasch, B., Büchel, C., Gais, S. & Born, J. Odor cues during slow-wave sleep prompt declarative memory consolidation. Science 315, 1426–1429 (2007).

77. Perl, O. et al. Odors enhance slow-wave activity in non-rapid eye movement sleep. Journal of Neurophysiology (2016).

78. Ngo, H.-V. V., Martinetz, T., Born, J. & Mölle, M. Auditory Closed-Loop Stimulation of the Sleep Slow Oscillation Enhances Memory. Neuron, 1–9 (2013).

79. Latchoumane, C. F. V., Ngo, H. V. V., Born, J. & Shin, H.S. Thalamic spindles promote memory formation during sleep through triple phase-locking of cortical, thalamic, and hippocampal rhythms. Neuron (2017).

80. Acsády, L, Kamondi, A, Sík, A, Freund, T & Buzsáki, G. GABAergic cells are the major postsynaptic targets of mossy fibers in the rat hippocampus. Journal of Neuroscience 18, 3386–403 (1998).

81. Shu, Y., Hasenstaub, A. & McCormick, D. A. Turning on and off recurrent balanced cortical activity. Nature 423, 288–293 (2003).

82. Timofeev, I., Grenier, F, Bazhenov, M, Sejnowski, T. J. & Steriade, M. Origin of slow cortical oscillations in deafferented cortical slabs. Cerebral Cortex 10, 1185–1199 (2000).

83. Sanchez-Vives, M. V. & McCormick, D. A. Cellular and network mechanisms of rhytmic recurrent activity in neocortex. Nature Neuroscience 3, 1027–1034 (2000).

84. Chauvette, S., Volgushev, M. & Timofeev, I. Origin of active states in local neocortical networks during slow sleep oscillation. Cerebral Cortex 20, 2660–2674 (2010).

85. Del Negro, C. A., Funk, G. D. & Feldman, J. L. Breathing matters. Nature Reviews Neuroscience 19, 351–367 (2018).

86. Vyazovskiy, V. V. et al. Local sleep in awake rats. Nature 472, 443–447 (2011).

87. Lemieux, M., Chauvette, S. & Timofeev, I. Neocortical inhibitory activities and long-range afferents contribute to the synchronous onset of silent states of the neocortical slow oscillation. Journal of Neurophysiology 113, 768–779 (2015).

88. Massimini, M., Huber, R., Ferrarelli, F., Hill, S. & Tononi, G. The Sleep Slow Oscillation as a Traveling Wave. Journal of Neuroscience 24, 6862–6870 (2004).

89. Carr, M. F., Jadhav, S. P. & Frank, L. M. Hippocampal replay in the awake state: A potential substrate for memory consolidation and retrieval. Nature Neuroscience 14, 147–153 (2011).

90. Jadhav, S. P., Kemere, C., German, P. W. & Frank, L. M. Awake hippocampal sharp-wave ripples support spatial memory. Science 336, 1454–1458 (2012).

## References

91. Paxinos, G., Franklin, K. B. J., Paxinos, G and Franklin, K., Paxinos, G. & Franklin, K. B. J. Mouse Brain in Stereotaxic Coordinates (2004).

92. Isosaka, T. et al. Htr2a-expressing cells in the central amygdala control the hierarchy between innate and learned fear. Cell 163, 1153–1164 (2015).

93. Lopes, G. et al. Bonsai: an event-based framework for processing and controlling data streams. Frontiers in Neuroinformatics 9 (2015).

94. Ferguson, J. E., Boldt, C. & Redish, A. D. Creating low-impedance tetrodes by electroplating with additives. Sensors and Actuators, A: Physical 156, 388–393 (2009).

95. Ludwig, K. A. et al. Poly(3,4-ethylenedioxythiophene) (PEDOT) polymer coatings facilitate smaller neural recording electrodes. Journal of Neural Engineering 8 (2011).

96. Hazan, L., Zugaro, M. & Buzsaki, G. Klusters, NeuroScope, NDManager: A free software suite for neurophysiological data processing and visualization. Journal of Neuroscience Methods 155, 207–216 (2006).

97. Mitra, P. P. & Pesaran, B. Analysis of dynamic brain imaging data. Biophysical Journal 76, 691–708 (1999).

98. Masimore, B., Kakalios, J. & Redish, A. D. Measuring fundamental frequencies in local field potentials. Journal of Neuroscience Methods (2004).

99. Barnett, L. & Seth, A. K. The MVGC multivariate Granger causality toolbox: A new approach to Granger-causal inference. Journal of Neuroscience Methods 223, 50–68 (2014).

100. Tort, A. B. L. et al. Dynamic cross-frequency couplings of local field potential oscillations in rat striatum and hip-pocampus during performance of a T-maze task. Proceedings of the National Academy of Sciences 105, 20517–20522 (2008).

101. Pettersen, K. H., Devor, A., Ulbert, I., Dale, A. M. & Einevoll, G. T. Current-source density estimation based on inversion of electrostatic forward solution: Effects of finite extent of neuronal activity and conductivity discontinuities. Journal of Neuroscience Methods 154, 116–133 (2006).

102. Ranck, J. B. Specific impedance of rabbit cerebral cortex. Experimental neurology 7, 144–152 (1963).

103. Logothetis, N. K., Kayser, C. & Oeltermann, A. In vivo measurement of cortical impedance spectrum in monkeys: implications for signal propagation. Neuron 55, 809–823 (2007).

104. Nicholson, C. & Freeman, J. A. Theory of current source-density analysis and determination of conductivity tensor for anuran cerebellum. Journal of Neurophysiology 38, 356–368 (1975).

105. Csicsvari, J. Massively parallel recording of unit and local field potentials with silicon-based electrodes. Journal of Neurophysiology 90, 1314–1323 (2003).

106. Buzsaki, G., Lai-Wo S. L. & Vanderwolf, C. H. Cellular bases of hippocampal EEG in the behaving rat. Brain Research Reviews 6, 139–171 (1983).

107. Ylinen, A. et al. Intracellular correlates of hippocampal theta rhythm in identified pyramidal cells, granule cells, and basket cells. Hippocampus 5, 78–90 (1995).

108. Senzai, Y. & Buzsaki, G. Physiological properties and behavioral correlates of hippocampal granule cells and mossy cells. Neuron 93, 691–e5 (2017).

109. Chung, J. E. et al. A fully automated approach to spike sorting. Neuron 95, 1381–1394.e6 (2017).

110. Csicsvari, J., Hirase, H., Czurko, A. & Buzsaki, G. Reliability and state dependence of pyramidal cell-interneuron synapses in the hippocampus: An ensemble approach in the behaving rat. Neuron 21, 179–189 (1998).

111. Freund, T. F. & Buzsaki, G. Interneurons of the hippocampus. Hippocampus 6, 347–470 (1996).

112. Henze, D. A. et al. Intracellular features predicted by extracellular recordings in the hippocampus in vivo. 84, 390–400 (2000).

113. DiCarlo, J. J., Lane, J. W., Hsiao, S. S. & Johnson, K. O. Marking microelectrode penetrations with fluorescent dyes. Journal of Neuroscience Methods 64, 75–81 (1996).

114. Summerlee, A. J. S., Paisley, A. C. & Goodall, C. L. A method for determining the position of chronically implanted platinum microwire electrodes. Journal of Neuroscience Methods 5, 7–11 (1982).

